# Monocyte-derived microglia with *Dnmt3a* mutation cause motor pathology in aging mice

**DOI:** 10.1101/2023.11.16.567402

**Authors:** Jung-Seok Kim, Sébastien Trzebanski, Sun-Hye Shin, Noa Chapal Ilani, Nathali Kaushansky, Marina Scheller, Aryeh Solomon, Zhaoyuan Liu, Oliver Aust, Sigalit Boura-Halfon, Lukas Amann, Marco Prinz, Florent Ginhoux, Roi Avraham, Carsten Müller-Tidow, Stefan Uderhardt, Ivan Milenkovic, Liran Shlush, Steffen Jung

**Author notes:** Corresponding author (S.J.).

## Abstract

Microglia are established in embryogenesis forming a self-containing cellular compartment resisting seeding with cells derived from adult definitive hematopoiesis. We report that monocyte-derived macrophages (MoMΦ) accumulate in the brain of aging mice with distinct topology, including the nigrostriatum and medulla, but not the frontal cortex. Parenchymal MoMΦ adopt *bona fide* microglia expression profiles. Unlike microglia, these monocyte-derived microglia (MoMg) are due to their hematopoietic origin targets of clonal hematopoiesis (CH). Using a chimeric transfer model, we show that hematopoietic expression of DNMT3A^R822H^, a prominent mutation in human CH, renders MoMg pathogenic promoting motor deficits resembling atypical Parkinsonian disorders. Collectively, these data establish in a mouse model that MoMg progressively seed the brains of aging healthy mice, accumulate in selected areas, and, when carrying a somatic mutation associated with CH, can contribute to brain pathology.

## Introduction

Microglia are critical players in central nervous system (CNS) development, homeostasis and immune defense ^1^. Microglia originate from primitive macrophage (MΦ) progenitors that arise in the yolk sac (YS), infiltrate the developing brain and clonally expand ^2,3^. YS-derived microglia persist throughout adulthood displaying considerable proliferation and self-renewal potential ^4,5^. Once definitive hematopoiesis commences in the mouse, hematopoietic stem cells (HSC) can seed the organism with MΦ that are disseminated via circulating blood monocytes. Monocyte-derived MΦ (MoMΦ) progressively replace embryonic MΦ in barrier tissues, such as the gut and skin, as well as selected other organs, ^6^. In the murine CNS, choroid plexus-associated MΦ have been reported to be replaced by MoMΦ ^7^. Other border-associated MΦ (BAM) in the perivascular ‘Virchow-Robin’ spaces and the leptomeninges are believed to remain under steady-state conditions largely of embryonic origin ^7^. Likewise, also parenchymal brain MΦ are thought to comprise during the life time of healthy mice exclusively YS-derived microglia and no MoMΦ ^8–10^. Moreover, even following challenges when monocytes infiltrate the brain to contribute activities distinct from microglia ^11^, they often only transiently seed the parenchyma ^12,13^, likely since MoMΦ fail to compete with microglia for to-be-defined niches in the CNS. Supporting this notion, once microglia are impaired by irradiation or genetic deficiencies, HSC-derived MΦ can efficiently and persistently colonize the brain parenchyma ^14–17^. Experimentally seeded parenchymal brain MoMΦ were shown to retain epigenomes and expression signatures distinct from YS-derived cells and respond differentially to challenges, even following prolonged brain residence ^14–17^. This includes, for instance, lack of expression of the transcriptional repressor Spalt Like Transcription Factor 1 (Sall1), a proposed hallmark of microglia identity ^14,18,19^.

Monocyte and MoMΦ contributions to brain pathologies are well documented in various animal models and the human disorders they represent. Involvement of these cells has been shown in the development, progression and resolution of experimental autoimmune encephalitis (EAE) ^20–22^, stroke ^23,24^, Alzheimer’s disease ^25,26^ and viral encephalitis ^27^. Whether and to what extend monocytes seed the healthy brain of mice, for instance with progressive aging, as recently suggested for humans ^28^, and whether monocytes give rise to persisting parenchymal MoMΦ and might thereby affect brain functions of the elderly is still under debate.

In contrast to microglia, MoMΦ are due to their derivation from HSC exposed to effects of somatic mutations that are associated with age-related clonal hematopoiesis (CH) ^29,30^. CH is defined by the expansion of hematopoietic and progenitor stem cells (HSPC) carrying leukemia – related mutations that accumulate with aging. CH-associated variants of the epigenetic writers and erasers Dnmt3a and Tet2, respectively, that affect the HSPC methylome have been shown to alter functions of monocyte-derived tissue MΦ and promote cardiovascular pathologies ^31–34^.

Here we took advantage of a fate mapping model to screen for the presence of monocyte-derived cells in the murine CNS. We corroborate the earlier notion that MoMΦ progressively replace BAM in the brain, but uncovered that, with increasing age, MoMΦ also accumulate in the parenchyma of selected brain regions of the healthy mice. Single-cell transcriptomics revealed that parenchymal MoMΦ in the aging brains adopt *bona fide* microglia signatures, while perivascular MoMΦ remain distinct from their embryo-derived counterpart. Finally, using a transplantation model, we show that monocyte-derived microglia (MoMg) that derive from HSC harboring a somatic mutation of the epigenetic writer DNMT3A, frequently found associated with human age-related CH, can cause a motor deficiency resembling atypical Parkinsonian disorders. Collectively, our results reveal that MoMg can progressively seed selected regions of the healthy mouse brain and present, when derived from hDNMT3A^R882H^ -expressing HSC, a risk factor for age-related brain pathologies.

## Results

### Monocyte-derived macrophages accumulate with age at the brain borders and in largely non-cortical parenchyma

Corroborating earlier observations ^35^, seminal fate mapping experiments in mice have established that microglia seed the developing brain in the embryo and subsequently maintain themselves through limited self-renewal ^3,36,37^. This has led to the prevalent notion that the parenchymal brain MΦ compartment of unchallenged mice does not receive input from adult hematopoiesis via blood monocytes, but comprises exclusively YS-derived microglia ^8–10^. In contrast, BAM were reported to be replaced to a various degree by MoMΦ^7^. To specifically screen the brain of healthy unmanipulated animals for monocyte-derived parenchymal and non-parenchymal MΦ, we took advantage of *Ms4a3^Cre^: R26-TdTomato* animals that introduce a permanent fluorescent label into monocytes, but not embryo-derived MΦ, due to *Ms4a3* expression in granulocyte monocyte progenitors (GMP) ^38^.

Flow cytometric analysis of brains of *Ms4a3^Cre^: R26-TdTomato* animals revealed the presence of TdTom^+^ cells, both in the leptomeninges and in the parenchyma (**Fig. 1a)**. The latter included, as expected ^7^ CD206^+^ BAM, but also TdTom^+^ cells expressing the microglia marker Tmem119, albeit at lower frequency. As earlier reported ^39^, parenchymal and leptomeningeal MΦ comprised populations that lacked and expressed surface MHCII respectively (**Suppl. Fig. 1a,b**). Percentages of MHCII^+^ cells were significantly higher in TdTom^+^ MoMΦ as compared to YS-derived MΦ (TdTom^−^) among both CD206^+^ and Tmem119^+^ brain MΦ.

**Figure 1.**
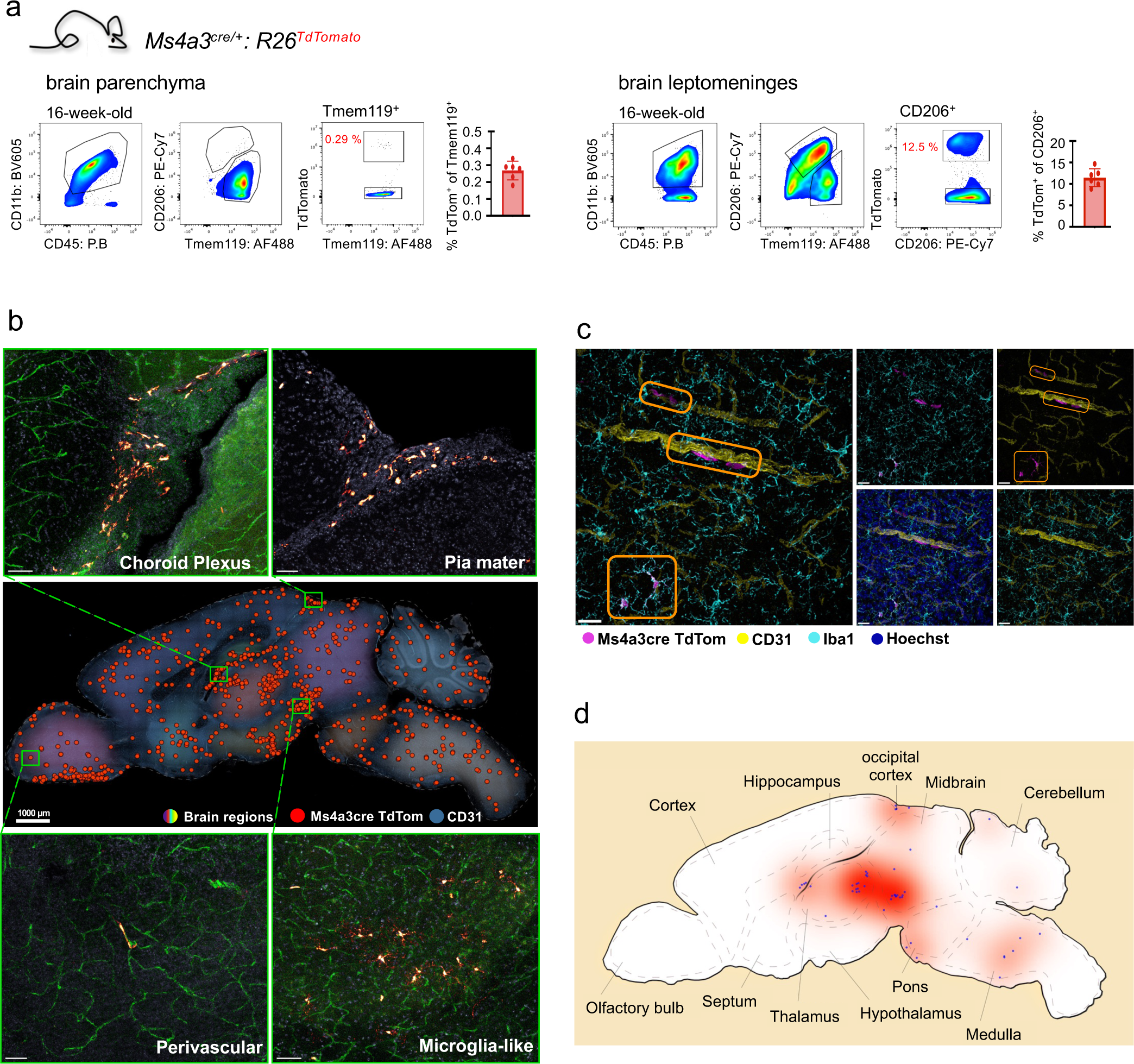
Flow cytometry and microscope images of brains from *Ms4a3^cre/+^: R26^TdTomato^* mice. (a) Representative flow cytometry results of parenchymal and leptomeningal cells from 16-week-old mouse brains. Pre-gated on CD45^+^ and Ly6C/G^−^. (n = 6). (b) Cryo-sagittal section of 6-month-old *Ms4a3^cre/+^: R26^TdTomato^* mouse brain stained with CD31 antibodies. Artificial colors indicate different brain regions, and red circles represent location of all TdTomato^+^ cells. Enlarged images come from the green rectangles in the sagittal section image. Scale is indicated in the sagittal section image, and scale bar of enlarged images is 100 µm. (c) Representative example image to describe how amoeboid perivascular cells were discriminated from ramified microglia-like cells among TdTomato^+^ population. Scale bar = 50 µm. (d) A schematic image to indicate the location of TdTomato^+^ microglia-like MoMΦ in the sagittal section of the brain shown in Fig 1B.

TdTom^+^ cells were also readily detected by immunofluorescence in sagittal thick sections of brains from *Ms4a3^Cre^: R26-TdTomato* mice (**Fig. 1b**). Parenchymal ramified Tmem119^+^ MoMΦ could be distinguished from vessel-associated amoeboid BAM and intravascular monocytes (**Fig. 1c, Suppl. Fig. 1c,d, Suppl. Movie 1**). Morphological discrimination revealed an accumulation of Iba1^+^ MoMΦ with microglia-like morphology in selected brain regions of the mice, including the occipital cortex, the nigrostriatal parenchyma and the medulla, but not the frontal and parietal-temporal cortex (**Fig. 1d**). The presence of ramified, microglia-like MoMΦ in the selected brain regions of aged animals was corroborated in an independent fate mapping model, i.e tamoxifen-treated *Cxcr4^CreER^: R26-TdTomato* mice, that displays a specific label of HSC-derived cells ^24^ (**Suppl. Fig. 1e,f**).

To explore a potential link between the observed MoMΦ in the CNS and aging, we performed a flow cytometry analysis of young and old mice. Specifically, we generated *Ms4a3^Cre^: R26-TdTomato: Cx3cr1^gfp^* double reporter (DR) animals that allow discrimination of TdTom^+^GFP^+^ MoMΦ from GFP^+^ YS-derived microglia and BAM (**Fig. 2a**). Blood analysis of DR mice confirmed that the vast majority of monocytes expressed both reporters, while granulocytes were as expected, TdTom^+^ GFP negative ^38,40^ (**Suppl. Fig. 2a**). Flow cytometry analysis of brain cells of DR mice allowed discrimination of GFP^+^ YS-derived cells and GFP^+^ TdTom^+^ MoMΦ among CNS border-associated CD206^+^ and parenchymal Tmem119^+^ MΦ populations (**Suppl. Fig. 2b,c**). As observed before, MHCII expression was largely restricted to MoMΦ (**Suppl. Fig. 2b,c**). Analysis of the parenchyma and the leptomeninges of young and old DR animals revealed that in both locations TdTom^+^ MoMΦ progressively accumulated with age (**Fig. 2b-d**). Flow cytometry analysis of brains of young and old single reporter animals corroborated this notion (**Suppl. Fig. 2d,e**). Histological analysis of sagittal brain sections of young and old *Ms4a3^Cre^: R26-TdTomato* mice confirmed the restricted localization of Iba1^+^ ramified MoMΦ in selected brain regions, excluding the frontal cortex (**Suppl. Table 1, Suppl. Fig. 2f**) and the increase of MoMΦ abundance with age (0.29% vs. 0.57 %) (**Fig. 2e**). Moreover, MoMΦ in aged brain were found clustered, suggesting clonal *in situ* expansion.

**Figure 2.**
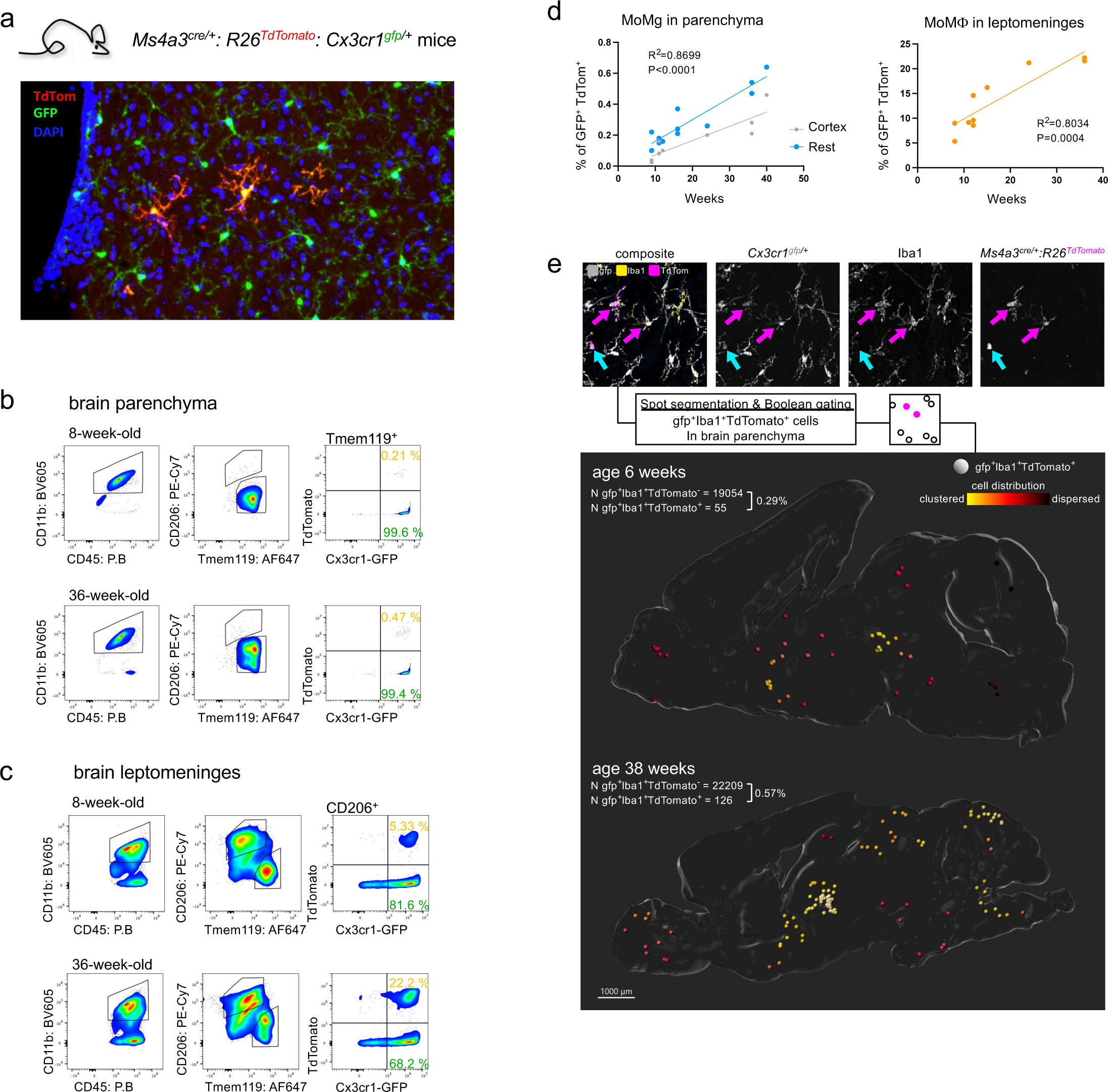
Flow cytometry and microscope images of brain from *Ms4a3^cre/+^: R26^TdTomato^: Cx3cr1^gfp/+^* mice. (a) A representative image of brain parenchyma from *Ms4a3^cre/+^: R26^TdTomato^: Cx3cr1^gfp/+^* mice. Sample was prepared as whole-mount section. GFP^+^ cells, YS-derived microglia; GFP^+^ TdTomato^+^ cells, MoMΦ. Scale is indicated in the image. (b) Representative flow cytometry result of brain parenchyma cells of 8- and 36-week-old mice. Pre-gated on CD45^+^ and Ly6C/G^−^. (c) Representative flow cytometry result of brain leptomeninges cells from 8- and 36-week-old mice. Pre-gated on CD45^+^ and Ly6C/G^−^. (d) Correlation analysis of MoMΦ and mouse ages in both parenchymal Tmem119^+^ and leptomeningeal CD206^+^ populations in *Ms4a3^cre/+^: R26^TdTomato^: Cx3cr1^gfp/+^* mouse brains (right graph, n = 12; left graph, n = 10). (e) Representative images of GFP and TdTomato signals, and Iba1 staining to discriminate GFP^+^ TdTomato^+^ MoMΦ in images (top), cryo-sagittal section of 6- and 38-week-old *Ms4a3^cre/+^: R26^TdTomato^: Cx3cr1^gfp/+^*mice (bottom). Magenta arrows indicate GFP^+^ TdTomato^+^ Iba1^+^ cells, and cyan arrow indicates GFP^+^ TdTomato^+^ Iba1^−^ cells. Microglia-like cells were identified based on the intensity of Iba1 staining and morphology. Percentage of GFP^+^ TdTomato^+^ MoMg is indicated in the images, and color scale indicates distance of adjacent cells. Scale is indicated in the image.

Immunofluorescence imaging analysis of the leptomeninges of DR mice confirmed abundant seeding of this BAM niche with MoMΦ, increasing with age (**Suppl. Fig. 2g)**. Of note, TdTom^+^ MoMΦ were more frequently found in the ventral leptomeninges, as compared to the dorsal side. This finding was corroborated by analysis of *Lyve1^ncre^: Cx3cr1^ccre^: R26-TdTom* mice ^41^. Also in these animals, the ventral surface of the brain harbored more MHCII^+^ and less Lyve1^+^ cells than the dorsal leptomeninges confirming regional difference of BAM replacement rates at the brain borders (**Suppl. Fig. 2h**).

Collectively, these data establish that the borders and the parenchyma of selected regions of brains of healthy aged mice are progressively seeded by monocyte-derived cells.

### Transcriptome comparison of YS-derived brain MΦ and MoMΦ accumulating in the aging mouse brain

Tissue MΦ identities are guided by environmental cues that the cells receive in their respective anatomic niches ^42^. Engrafted HSC-derived parenchymal brain MΦ have been shown to fail to adopt transcriptomes and epigenomes of microglia and remain distinct entities ^14–16^. This inability of the graft to establish a ‘microglia signature’ could be due to the irradiation and BM transfer. To investigate whether MoMΦ that arise in unmanipulated aging animals can acquire *bona fide* resident brain MΦ identities, we performed a comparative single-cell (sc) transcriptome analysis of MoMΦ and the respective original embryo-derived cells. Specifically, we sacrificed 1 year old DR animals and isolated Tmem119^+^ GFP^+^ microglia and Tmem119^+^ GFP^+^ TdTom^+^ parenchymal MoMΦ (MoMg), as well as CD206^+^ GFP^+^ leptomeningeal MΦ and CD206^+^ GFP^+^ TdTom^+^ leptomeningeal MoMΦ (leptoMoMΦ) from the brain leptomeninges followed by scRNA sequencing (**Fig. 3a**).

**Figure 3.**
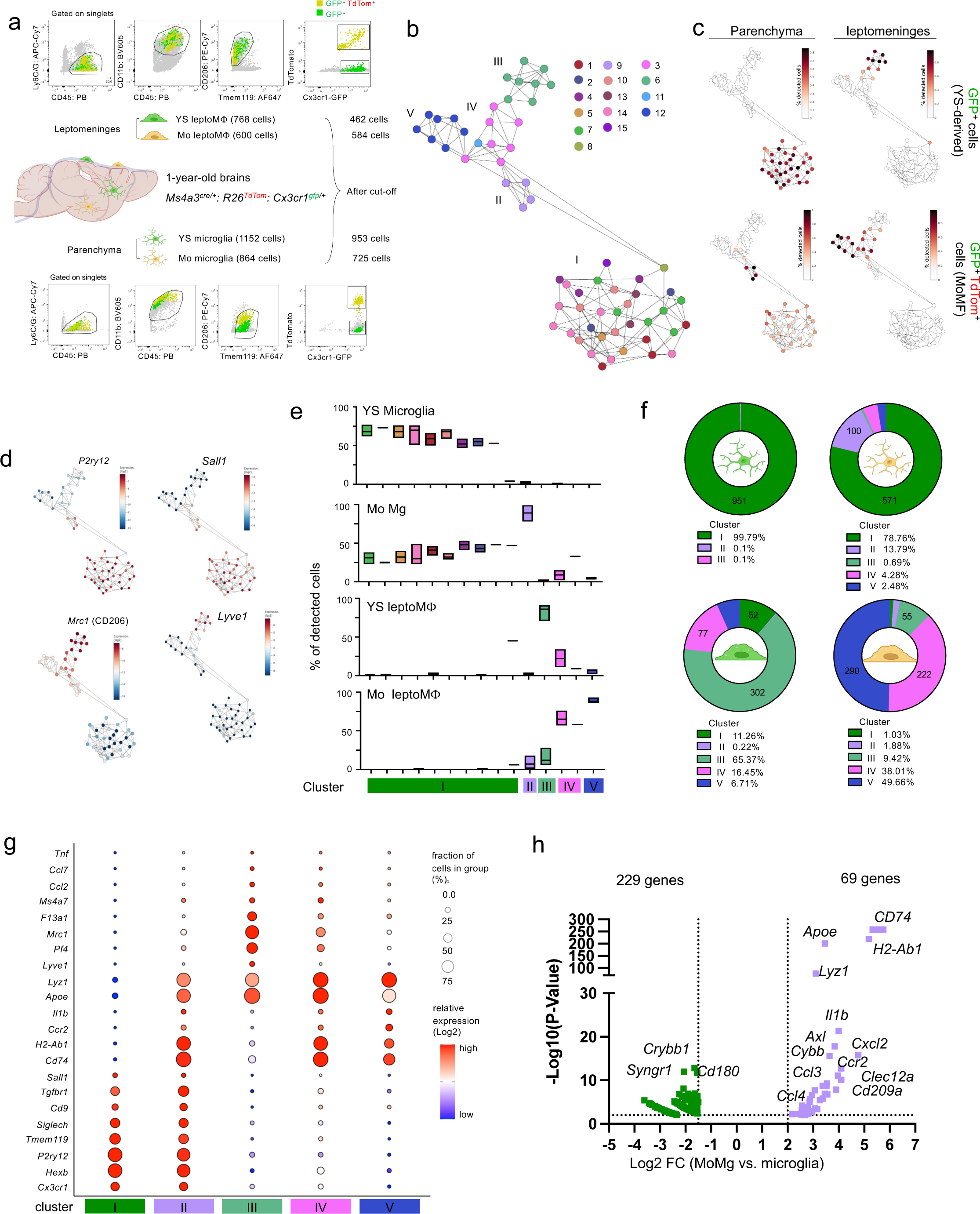
Single cell RNA sequencing of YS-derived MΦ and MoMΦ isolated from *Ms4a3^cre/+^: R26^TdTomato^: Cx3cr1^gfp/+^* parenchyma and leptomeninges. (a) Scheme and flow cytometry index sorting strategy for parenchyma and leptomeninges cells of *Ms4a3^cre/+^: R26^TdTomato^: Cx3cr1^gfp/+^* mouse brains. Yellow dots, GFP^+^ TdTomato^+^ cells; green dots, GFP^+^ cells. 5 brains from 1-year old mice were used for sorting, retrieved numbers are indicated in the figure. (b) MetaCell analysis identifying 15 cell types, following into 5 clusters (I-V). 11 cell types were located in the microglia and MoMg clusters (I, II), and 4 cell types were located in the leptoMΦ and leptoMoMΦ clusters (III-V). (c) MetaCell plots describing the correlation of index sorted cells and Meta cells. Color gradient indicates proportion of cells in four sorted subpopulations (parenchymal and leptomeningal GFP^+^ and GFP^+^ TdTomato^+^). (d) MetaCell plots showing *P2ry12, Sall1, Mrc1 (CD206),* and *Lyve1* gene expression. (e) Bar graphs showing the distribution and percentage of the sorted cells in each Meta cell types. Note that MoMg overlap with Meta cells referring to microglia. (f) Pie graphs showing how many sorted cells are placed in each cell cluster. (g) Dot plot showing representative gene expression level and fraction of positive cells in each cell cluster. (h) Volcano plots showing differentially expressed genes between microglia and MoMg in parenchyma. Only significantly different genes are presented, and dot colors indicate cell clusters. Significance means -Log10 (p value) > 1.5, Log 2 (fold change) > 1.

MetaCell analysis ^43^ grouped cells into five main clusters (**Fig. 3b**). Cluster I represented microglia and MoMg, as indicated through alignment with sorted Tmem119^+^ GFP^+^ microglia and Tmem119^+^ GFP^+^ TdTom^+^ parenchymal populations, and the expression of marker genes, such as *P2yr12* (**Fig. 3c,d, Suppl. Fig. 3a**). Cluster II was specific to MoMg (**Fig. 3c**). Cluster III - V referred to leptomeningeal cells, as indicated by *Mrc1* and *Pf4* expression, with cluster III comprising YS-leptoMΦ and Clusters IV and V representing leptoMoMΦ (**Fig. 3c,d, Suppl. Fig. 3b**). Highlighting their ability to adopt microglia identity, MoMg contributed to all 33 MetaCells covered by YS-derived microglia, although to varying degree (**Fig. 3e,f, Suppl. Fig. 3c**). MoMg in addition formed 4 MetaCells that had no microglia counterparts (cluster II), characterized by high expression of MHCII-related genes (*Cd74*, *H2ab-1*), as well as *Apoe* (**Fig. 3g,h, Suppl. Fig. 3d, e**). These MoMg-specific MetaCells harbored *Ccr2* transcripts, supporting the notion of their recent derivation from Ly6C^hi^ monocytes (**Fig. 3g,h, Suppl. Fig. 3f**). Surprisingly, and unlike what was reported for experimentally engrafted MΦ ^14–16^, MoMg, but not leptoMoMΦ, also expressed *Sall1* (**Fig. 3d,g**).

LeptoMoMΦ segregated into two clusters (IV, V) (**Fig. 3b**). As opposed to MoMg, LeptoMoMΦ did show only very minor overlap with their YS-derived LeptoMΦ counterpart (**Fig. 3e,f**). Higher expression of *Ccr2, Ly6c2* and lower expression of *Pf4, Mrc1* suggested that Cluster IV represents more recent recruits (**Fig. 3g, Suppl. Fig. 3g**), but both LeptoMoMΦ clusters were characterized by high expression of MHCII-related genes (**Fig. 3g, Suppl. Fig. 3g**). Notably, and in contrast to YS-leptoMΦ, leptoMoMΦ did not acquire Lyve1 expression in the tested time frame (**Fig. 3d**).

Taken together, these data establish that, as opposed to what was reported for experimentally engrafted cells ^14–16^, MoMg that accumulate in the aging brain of mice acquire transcriptomic identities of YS-derived microglia. In contrast, leptomeningeal monocyte-derived cells remained in the time frame analyzed distinct from YS-derived BAM.

### BM transplantation yields HSC-derived brain MΦ without monocytic intermediate and MoMΦ that accumulate with age

Following irradiation and BM transfer (BMT), brains of recipient mice are efficiently seeded with graft-derived MΦ that expand with age ^14^. Analysis of chimeras generated with *Ms4a3^Cre^: R26-TdTomato* BM (**Fig. 4a**) revealed that also MoMΦ generated in these mice accumulate first in the nigrostriatal regions, like MoMΦ that arise with age (**Fig. 4b, Suppl. Fig. 4 a,b**). This preferential seeding of selected brain regions was corroborated in BM chimeras lacking reporter gene insertions (**Suppl. Fig. 4c**).

**Figure 4.**
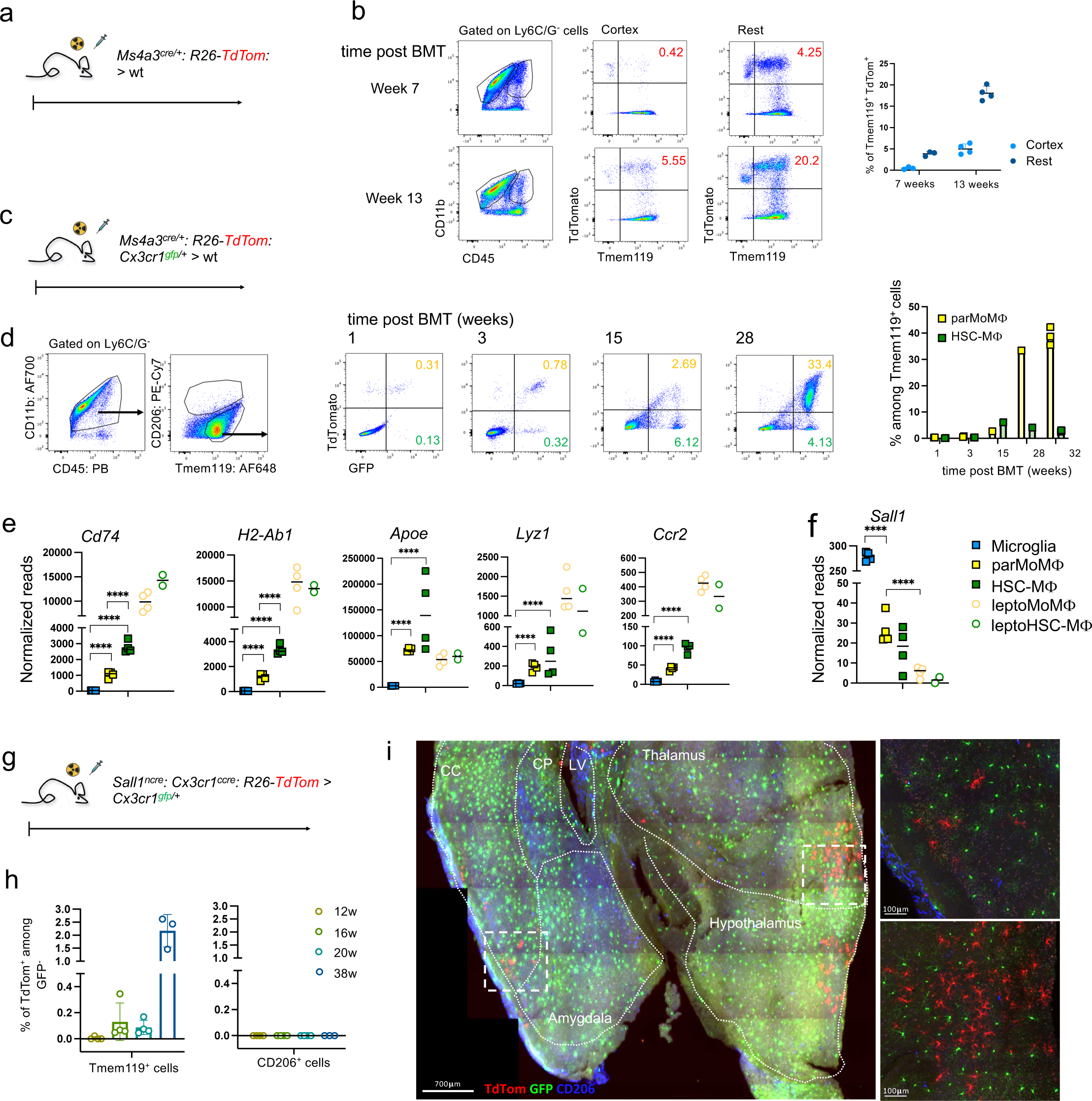
Flow cytometry and bulk RNA sequencing analysis of brain MΦ retrieved from BM chimeras. (a) BMT scheme with *Ms4a3^cre/+^: R26^TdTomato^* BM transferred to lethally irradiated WT mice. (b) Representative flow cytometry plots of parenchymal brain cells 7 and 13 weeks after BMT. Bar graph show percentages of Tmem119^+^ TdTomato^+^ cells in indicated brain regions at different time points. Flow cytometry plots are pre-gated on CD45^+^, Ly6C/G^−^. (7 weeks, n = 3; 13 weeks, n = 4). (c) BMT scheme with *Ms4a3^cre/+^: R26^TdTomato^: Cx3cr1^gfp/+^* BM transferred to lethal dose irradiated WT mice. (d) Representative flow cytometry plots of parenchymal brain cells pre-gated on CD45^+^, Ly6C/G^−^ 1 week, 3 weeks, 15 weeks and 28 weeks after BMT. Bar graph show percentages of engrafted cells among Tmem119^+^ population at different time points. (n = 3, per time point). (e) Representative gene expression level of different cell populations from bulk RNAseq with sorted cells of *Ms4a3^cre/+^: R26^TdTomato^: Cx3cr1^gfp/+^* BM transferred brain 32 weeks after BMT. (n = 4, except leptoHSC-MΦ n = 2) (f) *Sall1* mRNA expression levels in different cell populations from bulk RNAseq data. (n = 4, except leptoHSC-MΦ n = 2). (g) BMT scheme with *Sall1^ncre^: Cx3cr1^ccre^: R26^TdTomato^* BM transferred into irradiated *Cx3cr1^gfp/+^* mice. (h) Bar graph summarizing percentages of TdTomato^+^ cells among GFP^−^ Tmem119^+^ population in chimeras sacrificed at indicated time post BMT. (n = 3, in each time point) (i) Representative coronal whole-mount section image of *Sall1^ncre^: Cx3cr1^ccre^: R26^TdTomato^* > *Cx3cr1^gfp/+^* chimera brain at 38 weeks after BMT. Sample was prepared as section. Brain regions are indicated. Lower panel images are enlargement of rectangles on coronal section. Scale is indicated in each image. CC, cerebral cortex; CP, caudoputamen; LV, lateral ventricle.

BM grafts comprise HSC and myeloid precursors and fractionation experiments have suggested that engrafted Cx3cr1^−^ hematopoietic precursors directly enter the CNS and differentiate into MΦ that take long-term residence in the brain of chimeras ^13^. To define the origin of engrafted brain MΦ in irradiation chimeras and elucidate their relation to MoMΦ accumulating in aging brains (**Fig. 2d**), we generated chimeras using *Ms4a3^Cre^: R26-TdTomato: Cx3cr1^gfp^* (DR) BM (**Fig. 4c**). Specifically, in these DR-engrafted animals MoMΦ are GFP^+^ TdTom^+^ and can be discriminated from HSC- or early precursor-derived cells that might have skipped the GMP stage and hence remain GFP^+^ TdTom^−^ cells, here termed ‘HSC-MΦ’. Indeed, GFP^+^ TdTom^−^ HSC-MΦ could be readily detected in DR > wt chimeras, early after transfer (**Fig. 4d**). Time course analysis revealed however, that HSC-MΦ never reached more than ∼ 5% of the parenchymal brain MΦ population (**Fig. 4d**). In contrast, GFP^+^ TdTom^+^ MoMΦ developed only with time, but constituted by 8 months almost half of the Tmem119^+^ brain MΦ compartment.

To directly compare engrafted HSC- or early precursor-derived cells with engrafted MoMΦ we sorted the respective cells from brain parenchyma and leptomeninges of DR > wt chimeras, and performed a bulk RNAseq (**Suppl. Fig. 4d**). Heatmap analysis highlighted the distance of both MoMg and HSC-MΦ from YS-derived microglia corroborating an earlier study ^14^ (**Suppl. Fig. 4e-g**). Notably, MoMg and HSC-MΦ mostly differed in quantitative rather than qualitative expression of selected genes, including MHCII-related transcript (*Cd74, H2-Ab1*) (**Fig. 4e**).

Interestingly, MoMg and to a lesser extend HSC-MΦ displayed expression of *Sall1*, as compared to leptomeningeal MΦ (**Fig. 4f**), suggesting that also engrafted cells in BM chimeras can acquire this proposed ‘microglia signature’ marker ^14,18,19^. To independently probe whether engrafted MoMΦ can activate the *Sall1* locus we next generated chimeras with BM derived from *Sall1^ncre^: Cx3cr1^ccre^: R26-TdTom* mice, in which a binary transgenic splitCre system confers a permanent label to cells that co-expressed *Cx3cr1* and *Sall1* ^41^ (**Fig. 4g**). In line with our earlier report ^14^, up to three months after engraftment we did not detect any TdTom^+^ cells in the brain of these chimeras by flow cytometry (**Fig. 4h, Suppl. Fig. 4h**). Thereafter, however, the Tmem119^+^ parenchymal MΦ compartment, but not CD206^+^ BAM, displayed progressive accumulation of TdTom^+^ cells. In the old chimeras, labeled ramified microglia-like cells could also readily be detected by microscopy in distinct clusters, suggesting clonal expansion of the cells (**Fig. 4i**).

Collectively, these data establish that like age-related MoMg, engrafted MoMΦ accumulate in selected regions, that most engrafted CNS MΦ in brains of BM chimeras are monocyte-derived and that these cells approach with time microglia identity, as indicated by the acquisition Sall1 expression.

### Impact of hematopoietic hDNMT3A^R882H^ expression on the MoMg compartment

Unlike microglia, MoMΦ can carry somatic mutations acquired by proliferating HSC and resulting in age-related CH ^29,30^. Among the most common human CH variants are mutations affecting genes encoding DNA (cytosine-5)-methyltransferase 3A (DNMT3A) and Tet Methylcytosine Dioxygenase 2 (TET2) that add or remove DNA methylation (mC), respectively, and thereby modulate epigenetic gene regulation ^44–46^. Variants of these epigenetic modifiers predispose for hematological malignancies ^30,47^ and were shown to affect cardio-vasculature pathologies by altering expression profiles of monocyte-derived tissue MΦ ^31–33,48^.

To probe for the impact of a CH-associated somatic mutation on MoMg functions in the CNS and potential consequences on brain physiology, we took advantage of a mouse model that harbors a replacement of the murine *Dnmt3a* gene with a cDNA encoding the human DNMT3A^R882H^ variant ^49,50^. Upon introduction of a *Vav^Cre^* allele, HSC of these animals express hDNMT3A^R882H^ which alters, as reported for human cells ^51,52^ with time the HSPC methylome ^49^. To seed the CNS with MoMg derived from hDNMT3A^R882H^ HSC, we engrafted *Cx3cr1^gfp/+^* animals with BM of 10-month-old *Vav^Cre^: hDNMT3A^R822H^* mice, or age matched Cre-negative littermate BM for control (**Fig. 5a**). WT and mutant BM showed similar engraftment efficiency as indicated by blood and brain analysis for the presence of GFP-negative monocytes or MΦ (**Fig. 5b, Suppl. Fig 5a,b**). To gauge the impact of their derivation from hDNMT3A^R882H^ HSC, we isolated parenchymal and leptomeningal MoMΦ according to Tmem119, CD206 and GFP expression and performed bulk RNA sequencing. K-means clustering showed that microglia of the two types of BM chimeras displayed essentially identical transcriptomes, while control and mutant MoMg were distinct from host microglia (**Fig. 5c, Suppl. Fig. 5c**). Specifically, like control MoMg, MoMg that were derived from hDNMT3A^R882H^ HSC (‘hDNMT3A^R882H^ MoMg’) showed an ‘activation signature’ identified in the heatmap as cluster I-IV, spanning a total of 3153 DEG as compared to microglia. Cluster III, IV comprised 1966 genes which displayed higher expression in the mutant MoMg vs control MoMg, including MHCII-related genes (*Cd74, H2-Ab1)* and *Slc15a2*, encoding the proton-coupled oligopeptide transporter PepT2 (**Fig. 5d,e**). Notably, hDNMT3A^R882H^ MoMg also expressed significant levels of *Ccl5*, a chemokine that has been associated with brain pathologies and was shown to act on immune cells and neurons ^53–55^. Cluster I and II comprised 1187 genes expressed higher by control MoMg, than mutant MoMg, including *Plac8*. Also, mutant leptoMoMΦ displayed subtle changes as compared to control leptoMoMΦ, including elevated expression of *Slc15a2* and *Spint1* (**Suppl. Fig. 5d-f**). Of note, differences of mutant and control MoMΦ were more pronounced in the K-means analysis as compared to the Volcano plot. This is likely due to the fact that hypomethylation in HSPC of the hDNMT3A^R882H^ chimeras has a global effect on the expression profiles of their progeny and MoMΦ therefore are likely transcriptionally heterogeneous.

**Figure 5.**
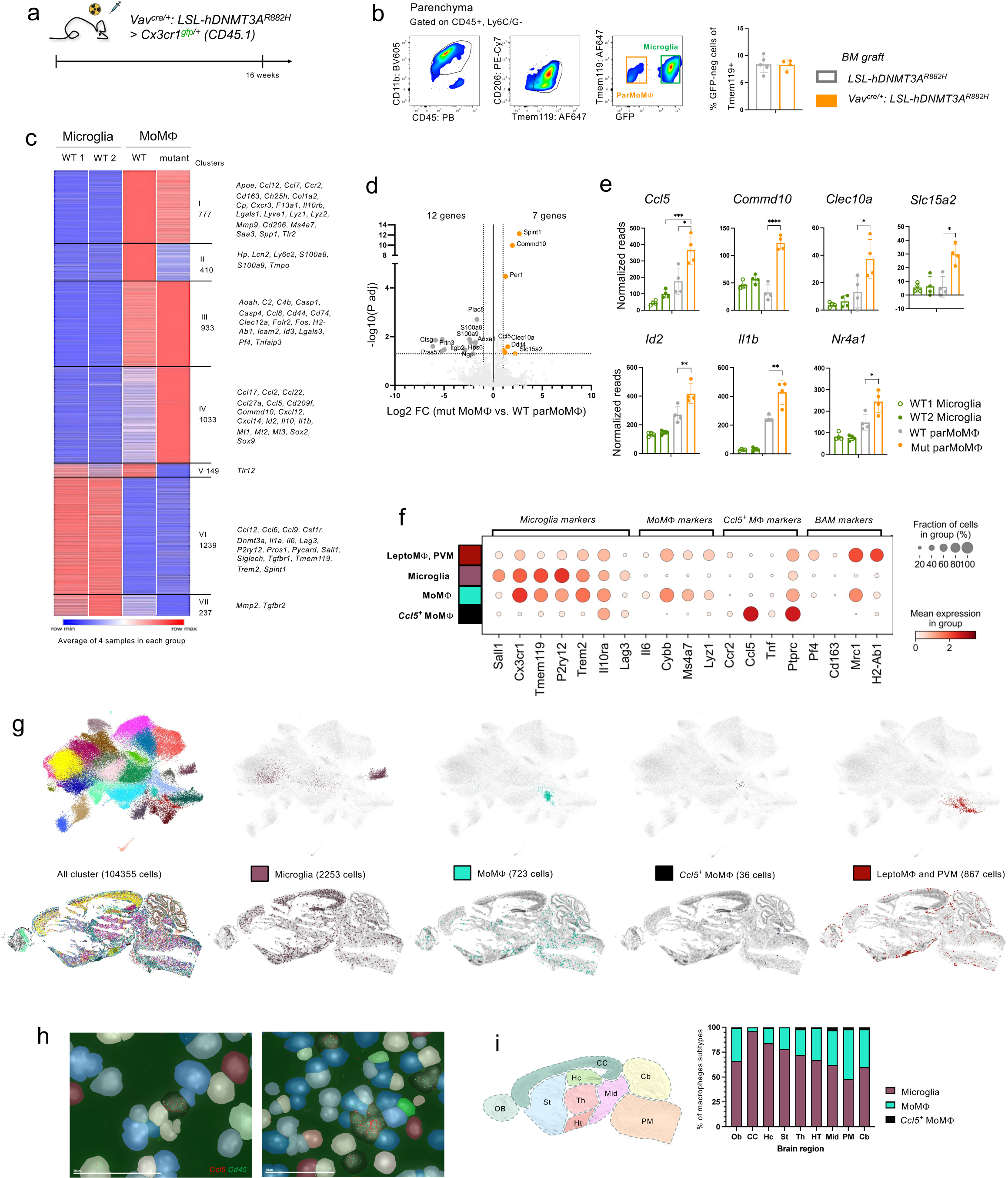
Flow cytometry, bulk RNA sequencing, and spatial transcriptomic results of *Vav^cre/+^: hDNMT3A^R822H/+^* BM transfer experiment. (a) BMT scheme of *Vav^cre/+^: hDNMT3A^R822H/+^* (and *hDNMT3A^R822H/+^* (control)) BM transferred into lethally irradiated *Cx3cr1^gfp/+^*mice. (b) Representative flow cytometry plots of brain parenchymal cell preparations used for cell sorting. Cells are pre-gated for CD45, Ly6C/G. 18 weeks after BMT and a Bar graph shows chimerism of Tmem119^+^ brain cells in *Vav^cre/+^: hDNMT3A^R822H/+^* and control chimeras (n = 5, wt BM, n = 4, mutant BM). (c) Heatmap of bulk RNA sequencing result. Average values of 4 samples, each. (d) Volcano plot comparison of control and mutated MoMΦ in brain parenchyma. Significant DEG are indicated with bold dots. Significance means -Log10 (adj. p-value) > 1.5, Log 2 (fold change) > 1. (e) Representative DEG between engrafted cells retrieved from control BM and mutant BM recipients. (n = 4). (f) Spatial transcriptomic data showing expression levels of selected genes in microglia, MoMg, *Ccl5^+^* MoMg and leptomeningeal MΦ/ PVM clusters. (g) Spatial transcriptomic data analysis, UMAP (top) and spatial distribution (bottom) of all cell clusters across sagittal brain section of *Vav^cre/+^: hDNMT3A^R822H/+^* BM-transplanted *Cx3cr1^gfp/+^* mouse (18 weeks after transplantation). Microglia, MoMg, *Ccl5*^+^ MoMg, and leptomeningeal MΦ / PVM clusters are presented separately. (h) Representative images of *Ccl5* and *CD45* expression in *Ccl5^+^* MoMg. (i) Schematic of brain regions and bar graph summarizing distribution of cell types in different brain regions of chimera. Cb, cerebellum; CC, cerebral cortex; Hc, hippocampus; HT, hypothalamus; Mid, mid brain; OB, olfactory bulb; PM, pons and medulla; St, striatum; Th, thalamus.

To locate MoMΦ in the brain tissue context we performed spatial transcriptomics using the Merscope platform (Vizgen). Specifically, we used a custom list comprising 140 genes to analyze sagittal brain sections of *Vav^Cre^: hDNMT3A^R822H^* > Wt BM chimeras (**Suppl. Data 1**). Host microglia and BAM could be discriminated according to co-expression of key markers such as *P2yr12* / Tmem119, and *Pf4 / Lyve1*, respectively (**Fig. 5f)**, and displayed discrete anatomic distribution (**Fig. 5g, Suppl. Movie 2**). MoMg comprised two populations, one of which displayed *Ccl5* expression (**Fig. 5f-h**). Interestingly, and in line with the imaging data (**Fig. 1d, Suppl. Fig. 4a**), MoMg were excluded from frontal cortical areas but found diffusely spread in nigrostriatal and medullary area (**Fig. 5i**).

Collectively, these data establish that the hDNMT3A^R882H^ -related epigenome alteration impacts the transcriptome of MoMΦ in the brain of the recipient animals, including the expression of Ccl5.

### Mice harboring hDNMT3A^R882H^ MoMg develop a motor disability

To investigate whether hDNMT3A^R882H^ MoMg impact brain functions we subjected the chimeras that received *Vav^Cre^: hDNMT3A^R822H^* BM or *hDNMT3A^R822H^* control BM to a marker-less CatWalk gait analysis which records paw-contacts together with body silhouettes ^56^. Principle component analysis (PCA) of the collected gait parameters, two, three, and four months after engraftment, revealed segregation of the recipients of the mutant BM that developed in particular at the late time point (**Fig. 6a, Suppl. Fig. 6a**). Specifically, as compared to controls, *Vav^Cre^: hDNMT3A^R822H^* BM recipients displayed reduced average speed and cadence, as well as extended standing on three and four legs (**Fig. 6b, Suppl. Fig. 6b, Suppl. Data 2**). As compared to controls, chimeras harboring mutant MoMg also performed less well in a pole test, a widely used assay for basal ganglia-related movement disorders in mice ^57^ (**Fig. 6c**).

**Figure 6.**
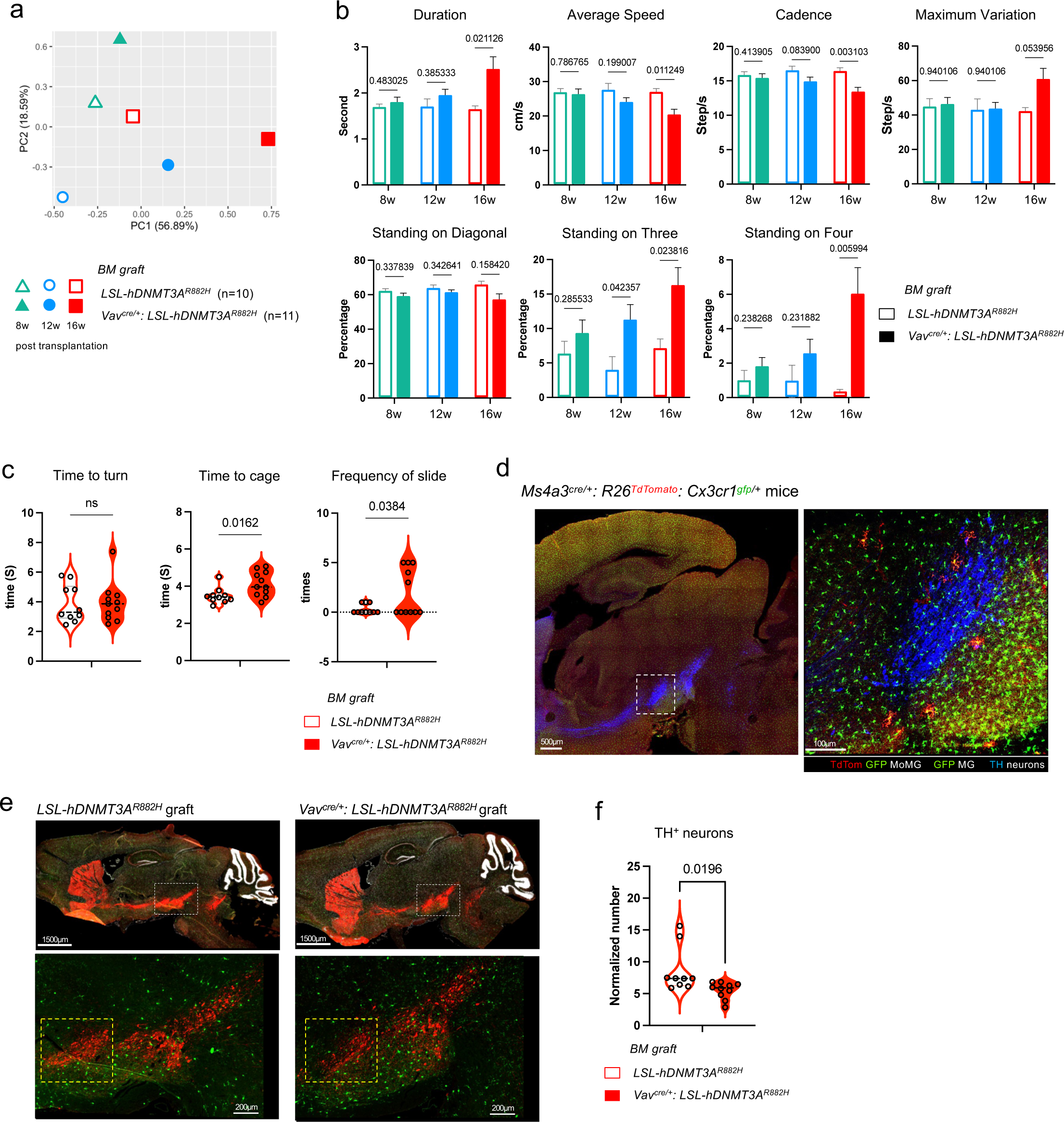
Motoric impairment caused by *hDNMT3A^R822H/+^* MoMg. (a) PCA plots of 120 parameters acquired with CatWalk gait analysis system. Time points of CatWalk analysis is 8, 12 and 16 weeks after BMT. (n=10, per group) (b) CatWalk gait results of parameters that indicated motor impairment of mutated BM recipients compared with the control BM recipients. Multiple T-test was performed for statistical comparison, and specific q values are written in the graphs. (n = 10, per group). (c) Pole task performed with recipients of mutant and control BM recipients. Task has performed 16 weeks after BMT. Time to turn, time to cage and frequency of slide during the task were measured manually. Means of three times trails are presented in the graphs for each mouse. P-values after t test are indicated in each plot. (n = 10). (d) Representative immunohistochemical analysis of sagittal brain section of *Ms4a3^cre/+^: R26^TdTomato^: Cx3cr1^gfp/+^*mouse indicating presence of GFP^+^ TdTomato^+^ MoMg in TH^+^ substantia nigra pars compacta. (e) Representative immunohistochemical analysis of sagittal brain section of *LSL-hDNMT3A^R882H^* or *Vav^cre/+^: LSL-hDNMT3A^R882H^* BM transferred mouse. Stained with anti-TH antibodies. Lower panel is enlargement image of upper panel respectively. (f) Quantification of TH^+^ neurons in the substantia nigra pars compacta of brains of mutant and control BM recipients. Data are based on the analysis of sections of 5 animals per group, 2 sections each. For details of the area definition and calculation see Supplementary Figure 6C, D.

The progressive gait impairment and poor performance in the pole test of the *Vav^Cre^: hDNMT3A^R822H^* BM recipients was reminiscent of a hypokinetic motor phenotype-associated with parkinsonism ^58^ that results from dopaminergic deficiencies. Analysis of brains of DR animals, showed the presence of MoMΦ in the *substantia nigra pars compacta* (SNc), delineated by tyrosine hydroxylase (TH) staining, and established as a control center for motor performance (**Fig. 6d**). To investigate if hDNMT3A^R882H^ MoMg affect dopaminergic circuits we analyzed the abundance of TH^+^ neurons in this brain area of the two groups of chimeras. Immuno-histochemical quantification / TH^+^ volume revealed a loss of TH^+^ neurons in mice harboring hDNMT3A^R882H^ MoMg, as compared to controls (**Fig. 6e,f, Suppl. Fig. 6c,d**), suggesting local impairment of neuronal transmission.

Collectively, these data establish that MoMg accumulate in the aging brain in nigrostriatal regions and when derived from hDNMT3A^R882H^ HSC affect dopaminergic circuits and cause with time a motoric deficiency.

## Discussion

Here we establish that monocyte-derived MΦ progressively seed specific selected regions of the brain in healthy aging mice, excluding large areas of the cortex. Unlike leptomeningal MoMΦ, parenchymal MoMg that accumulate with age adopt expression signatures of *bona fide* YS-derived microglia. Unlike microglia, MoMg are however subject to somatic mutations associated with CH, and as established here in a mouse model for the DNMT3A^R822H^ variant, MoMg can hence constitute an age-related risk factor for brain pathologies.

Monocyte-derived brain MΦ have been reported earlier, but these studies were largely focused on conditions of experimental challenge ^20–25,59^. This includes a recent report by the Ginhoux group involving the *Msa4a3^Cre^* fate mapping model used in this study ^60^. Here we establish that MoMΦ seed the brain of aging animals also in absence of challenge with a particular topology restricted to specific brain regions, albeit at low numbers. Arguably, mice maintained under hyper-hygienic specific pathogen free (SPF) housing conditions are, as compared to animals roaming in the wild, likely to show low MoMg seeding rates, both in the parenchyma and the CNS borders. Likewise, MoMg contributions to the CNS MΦ compartments can be predicted to more substantial in humans and could well increase following concussions and other parenchymal traumata. In line with this notion, a recent study reported a high frequency of MoMg in brains of octogenarians ^28^, although the medical history of these individuals remains unknown.

Using fate mapping we show that MoMg accumulate with age in various brain regions, with the notable exception of the frontal cortex. The reason for this selected MoMg seeding pattern, which is also observe in aging irradiation chimeras, remains to be investigated. Differential seeding could be due to regional differences in the microvasculature or the BBB ^61,62^. Increased vessel density or a more permissible barrier could allow increased entry of blood monocytes from the circulation into the parenchyma. Alternatively, but not mutually exclusive, vasculature leak could result in enhanced exposure of local microglia to environmental and serum factors, such as endotoxins and fibrinogen ^63–65^. Collectively, this could lead to repeated regional microglia activation and eventual exhaustion of the cells resulting in impaired competition of microglia with MoMg. Of note, the available reports of monocyte-derived microglia in humans, as defined by CH mutations, so far refer to brain regions that overlap with the areas in which we observe MoMg accumulation in the aging mice, including the occipital cortex, medulla and putamen ^28,66^.

We show by scRNAseq analysis that MoMg that accumulate in aging brains acquire *bona fide* microglia transcriptomes. This included acquisition of Sall1 expression, a marker we and others had previously assumed to be restricted to YS-derived cells ^14–16,18^. Indeed, over time engrafted MoMg also acquire Sall1 expression. This is in line with recent data reported for human monocyte-derived microglia which were shown to harbor ATACseq signals in the Sall1 loci ^28^. These data suggest that, despite of their distinct origin, MoMg become with time identical to microglia on the transcriptome level. Future studies involving specific challenges will be required to establish if YS-Mg and MoMg are also functionally equivalent.

In aging unmanipulated mice, MoMg progressively seed largely non-cortical brain areas. This finding could explain reports of regional ‘microglia heterogeneity’. Specifically, MHC II expressing ‘microglial cells’ identified in the nigrostriatal compartment ^67^ could, for instance, represent MoMg, that as we show express MHC II. Moreover, also reported age-associated ventral tegmental area (VTA) - and SNc - specific alterations could be related to MoMg replacement of microglia ^68^.

Our study largely focused on parenchymal MΦ, but analysis of the leptomeninges also showed robust seeding of the BAM compartment with MoMΦ, in particular for the ventral sides of the aging brain. The BAM replacement rates we observed considerably exceeded earlier reports ^7,69^, which could be related to animal housing conditions but requires further studies. Notably, the scRNAseq analysis revealed that leptomeningeal MoMΦ that accumulate at the CNS borders retain transcriptomes distinct from their YS-derived counterpart.

Clonal hematopoiesis (CH) is defined by the acquisition of somatic mutations in HSC and occurs in up to one third of individuals aged > 60 years. CH prominently involves variants of epigenetic regulatory genes, such as DNMT3A and TET2, which constitute an increased risk for hematologic malignancies, cardiovascular events and stroke ^34,45,70,71^. Here we show in a mouse model that as a result of age-related seeding of selected brain regions with MoMg, the most common human CH mutation, DNMT3A^R822H^, presents a risk factor for motor pathology, Specifically, using a BM transfer model, we show that MoMg that derive from HSC that express the DNMT3A^R822H^ variant cause a motor impairment reminiscent of the hypokinetic motor phenotype seen in parkinsonism ^72^. In line with a Parkinson-related pathology, the presence of mutant MoMg in the *substantia nigra pars compacta*, resulted in a loss of dopaminergic TH^+^ neurons. Using spatial transcriptomics we show that mutant MoMg, including cells that express *Ccl5* are located in basal ganglia regions, in line with the possibility that they might directly affect the dopaminergic circuits. However, and as in individuals harboring CH variants, mutant monocytes in the chimeras also seed peripheral MΦ compartments, and we cannot rule out that this contributes to the observed phenotype.

We assume that the effect of the DNMT3A^R822H^ variant results from the altered HSC methylome of *Vav^Cre^: hDNMT3A^R822H^* mice ^49^ rather than the expression of the mutant epigenome writer in MoMΦ. Targeted expression of *hDNMT3A^R822H^*in microglia could be used to gauge a possible direct impact on mature MΦ, although this experiment is arguably artificial since microglia are not subjected to CH.

Interestingly and in line with our findings of the spatially restricted CNS seeding by MoMΦ, an unbiased screen for somatic mutation in human brain sections revealed cells harboring DNMT3A and TET2 variants restricted to the medulla, suggesting that these regions harbor islands of MoMg bearing CH mutations ^66^. The presence of CH mutations served also as a proxy to identify HSC-derived cells among the brain MΦ compartment in the recent study by Jaiswal and colleagues ^28^. The single-nucleus chromatin profiling of brain cells of octogenarian CH carriers revealed frequencies of MoMg considerably higher than what would be anticipated from fate mapping studies in mice, including this study. The difference of the MoMg abundance between the analyzed human and mice brains could be related to the life span difference of the analyzed organisms and the related differential exposure to environmental factors or pathogens, which remains unknown for the human individuals tested, but clearly warrants further dedicated study.

Our data in the mouse model suggest that MoMΦ that seed the nigrostriatal regions of aging brains impair dopaminergic circuits causing motor symptoms of parkinsonism. MΦ have been previously implicated in human PD-like pathology. Specifically, a common genetic cause of PD are mutations of the leucine rich repeat kinase 2 (LRRK2) gene ^73^ which is predominantly expressed in the CNS by MΦ. The molecular mechanism underlying the pathology we observe remains however to be defined. Emerging evidence suggests that cardiovascular pathologies and atherosclerosis exacerbated by CH mutation-afflicted MoMΦ involve distinct effector molecules, such as Il1β and IL6 ^31,74^. MoMg in the brain parenchyma, and even more so mutant MoMg, did express Il1β, but no IL6. Interestingly though, in particular the MoMg derived from hDNMT3A^R882H^ HSC also prominently expressed *Ccl5*, encoding a chemokine that acts on immune cells and neurons, and has been associated with brain pathologies, including PD ^53–55^. Definition of the functional relevance of specific circuits in the hDNMT3A^R882H^-driven motor disorder could be addressed by future MoMg-specific conditional mutagenesis in mice. Establishment of a link between CH, and in particular the hDNMT3A^R882H^ variant, with Parkinsonism or other CNS disorders will require dedicated and comprehensive studies on relevant human cohorts.

Our data show that in a mouse model, due to their hematopoietic origin, monocyte-derived brain macrophages can constitute a risk factor. However, the integration of MoMg into the aging brain microglia compartment and their adoption of *bona fide* microglia signatures, could well result in beneficial rejuvenation. Likewise, emerging data suggest that MoMΦ could counter AD pathogenesis ^26^.

Taken together, we establish that the parenchymal CNS MΦ compartment of unmanipulated aging mice not only comprises YS-derived microglia, but also monocyte-derived cells, that progressively seed selected largely non-cortical brain regions. With the example of MoMg derived from hDNMT3A^R882H^ expressing HSC, we demonstrate in a mouse model that MoMg can present a risk factor for age-related brain pathology, since they are, unlike microglia, CH targets. Together with a recent report ^28^ our study hence calls for further exploration a possible linkage of CH and neurological disorders in the elderly.

## List of Supplementary Materials

**Supplementary Data 1**. Gene list for spatial transcriptome analysis

**Supplementary Data 2**. Data of CatWalk analysis

**Supplementary Movie 1**. Cryo-sagittal section of *Ms4a3^cre/+^: R26^TdTomato^* mouse brain

**Supplementary Movie 2**. Spatial transcriptomic analysis of sagittal brain sections

**Supplementary Table 1**.

**Source File** (to be added upon acceptance)

## Data deposit

Data related to specific figures can be found in the Online Source file. Gene expression data sets are deposited at GEO (Accession # xxxx) (to be added upon acceptance)

## Author contributions

J-S. K. made the initial observation and conceived the project with S. J. S. T. S.-H. S. and S. B. H. helped with the analysis. N. K. and L. S. advised on CH. I. M. advised on brain histology and PD pathology. L. A. and M. P. provided tissue samples, A. S. and R. A. provided help with scRNAseq, and N. C. I. ran the MetaCell analysis for scRNAseq data. O. A. and S. U. performed imaging analysis. Z. L., F.G., M. S., C. M.-T. provided transgenic animals. J.-S. K. and S. J. wrote the manuscript.

## Acknowledgements

We would like to thank all members of the Jung laboratory, for advice and helpful discussion. We thank the Shahar lab for providing chimeras and the staff of the Weizmann Animal and flow cytometry facilities, as well as M. Tsoory for help with the CatWalk Gait analysis. S. J. is the incumbent of the Henry HG. Drake Professional Chair of Immunology. This project was funded by the Deutsche Forschungsgemeinschaft (DFG, German Research Foundation) – Project-ID 259373024 – TRR 167, a research grant from the Sagol Institute for Longevity Research and the Israeli Science Foundation (ISF) (grant # 696/21). S. U. is supported by the DFG (project-IDs 448121430, 405969122, 505539112), the Hightech Agenda Bavaria and an ERC starting grant (project-ID 101039438). Whole brain section imaging was performed on a DFG-funded confocal microscope (project-ID 261193037).

## Materials and Methods

### Animals

Mice used in this study were all on C57BL/6 background, including wild type animals, C57BL/6J-*Ms4a3^em2(cre)Fgnx^*/J mice (*Ms4a3^cre^*) (Jackson Laboratory Stock #036382) ^38^, B6.129P2(Cg)-*Cx3cr1^tm1Litt^*/J mice (*Cx3cr1^gfp^*) (Jackson Laboratory Stock # 005582) mice ^40^, B6.Cg-*Gt(ROSA)26Sor^tm9(CAG–tdTomato)Hze^*/J mice (Jackson Laboratory Stock # 007909) ^75^, C57BL/6 *Sall1^em1(Ncre)Jung^ Cx3cr1^em1(Ccre)Jung/^*J mice (*Sall1^ncre^:Cx3cr1^ccre^*) (Jackson Laboratory Stock #033318) ^41^, *B6.Cg-Tg(VAV1-cre)1Graf/MdfJ* mice (Vav-Cre) (Jackson Laboratory Stock #035670) ^76^, and LSL-*DNMT3A*^R882H^ mice ^49^. All *Ms4a3^cre^*, *Cx3cr1^gfp^* and *DNMT3A*^R882H^ animals in this study were heterozygotes. To generate BM chimeras, wild type C57BL/6 J mice (Harlan), *Cx3cr1^gfp^* mice or *B6.SJL-Ptprc^a^ Pepc^b^/BoyJ* (CD45.1) mice were used as recipient, and *Ms4a3^cre^: R26-TdTom^fl/fl^*, *Ms4a3^cre^: R26-TdTom^fl/fl^: Cx3cr1^gfp^*, Sall1^ncre^: Cx3cr1^ccre^: R26-TdTom^fl/fl^, *DNMT3A*^R882H^, *Vav^cre^*: *DNMT3A*^R882H^ mice, or wild type C57BL/6 J mice (Harlan) were used as donors. Recipient mice were lethally irradiated with a single dose of 950 cGy generated by XRAD 320 machine (Precision X-Ray (PXI) and BM was transferred the next day (5 × 10^6^ donor cells per mouse introduced via i.v. injection). All animals were maintained in a specific pathogen-free (SPF) facility with chow and water provided *ad libitum* and handled according to protocols approved by the Weizmann Institute Animal Care Committee as per international guidelines.

### Cell isolation from brain leptomeninges and parenchyma for flow cytometry

Mice were anesthetized with Pental (1:1 in PBS), and perfused with cold PBS. Whole mouse brains were isolated in intact condition, and incubated for 40 min at 37 °C with gentle shaking in the digestion solution (3 ml HBSS /per brain containing 2% BSA, 5 mM EDTA, and 0.2 mg/ml Collagenase type 2 (Worthington)). After the incubation leptomeningeal cells were washed out from the surface of mouse brains with pipetting of digestion solution on the surface, and collected in tubes with cold PBS after 100-µm mesh filter. Collected cells were centrifuged at 400 g at 4 °C for 5 min, and re-suspended with cold FACS buffer containing 2% FBS, 1 mM EDTA based on PBS w/o Ca_2_^+^ and Mg_2_^+^ for antibody staining. Intact brains were further digested for parenchymal cell isolation. Brains were dissected in the 1 ml of digestion solution containing 2% BSA, 1 mg/ml Collagenase D (Sigma) and 1 mg/ml DNase1 (Sigma), and incubated for 20 min at 37 °C. In the middle of incubation, brains were homogenized by pipetting. After the incubation the homogenate was filtered through a 100-µm mesh, and centrifuged at 970 g at 4 °C for 5 min. To enrich for brain cells, the cell pellet was re-suspended with 3 ml of 40% Percoll solution and centrifuged at 970 g, w/o acceleration and brake, at room temperature for 15 min. After the centrifugation, the myelin layer was removed and the rest of supernatant and cell pellets were washed in cold PBS. Cells were re-suspended with cold FACS buffer. For the blood analysis, blood was collected in a tube with heparin, and centrifuged at 400 g at 4 °C for 5 min. Blood cell pellets were re-suspended in 1ml ACK lysis buffer and incubated in room temperature for 5 min, followed by adding 10 ml cold PBS. After centrifugation, cell pellets were re-suspended with FACS buffer. Cells were kept on ice during all steps, except for the enzymatic digestion and cells for bulk-RNAseq and scRNAseq were processed always with actinomycin D (ActD) to prevent transcriptomic changes, as reported before ^39^.

### Flow cytometry and cell sorting

Cells were stained with antibodies against murine CD45 (30-F11, Biolegend cat. no. 103112, RRID: AB_312976), CD45.1 (A20, Biolegend cat. no. 110721, RRID: AB_492867), CD45.2 (104, Biolegend cat. no. 109829, RRID: AB_1186109), CD11b (M1/70, Biolegend cat. no. 101205, RRID: AB_312788), Ly6C/G (Gr-1) (RB6-8C5, Biolegend cat. no. 108411, RRID: AB_313376), I-Ab (AF6-120.1, Biolegend cat. no. 116421, RRID: AB_10613291), Tmem119(106-6, Abcam cat. no. ab225494, RRID: AB_2744673), CD206 (C068C2, Biolegend cat. no. 141720, RRID: AB_2562248), CD115 (AFS98, Biolegend cat. no. 135509, RRID: AB_2085222), Ly6C (HK1.4, Biolegend cat. no. 128015, RRID: AB_1732087). Methods adhered to published guidelines ^77^. Analysis was performed on a LSR Fortessa 4 lasers (BD Biosciences) or Aurora (Cytek Biosciences), and data were acquired with FACSDiva (BD Biosciences) and SpectroFlo (Cytek Biosciences). Post-acquisition analysis was performed using FlowJo software (Tree star, FlowJo, LLC). Bulk RNA sequencing and scRNA sequencing samples were flow sorted in 1.5ml tubes, and single cells were sorted in 384-well plate with a FACSAriaIII cell sorter (BD Bioscience) and BD FACSDiva software.

### Immunofluorescence microscopy

Mice were anesthetized with Pental (1:1 with PBS) and perfused with PBS. Brain tissues were excised and fixed for 2 hours in 4% paraformaldehyde in room temperature. Whole-mount prefixed tissues were blocked in 2% horse serum for 2 hours in room temperature and incubated with primary antibody overnight at 4 °C. After the incubation, tissues were washed three time in PBS and incubated with secondary antibody for 2 hours at room temperature. Tissues were incubated for 5 min with DAPI (1:10,000) (Sigma), and washed three time with PBS. For the cryosection, fixed brain tissues were incubated in 30% sucrose for 72 hours at 4 °C before optimal-cutting-temperature compound (TissueTek) imbedding and freezing. The brains were then sectioned with 60 µm thickness on slides. For the staining, the sections were blocked with 0.5% Triton X-100 in PBS containing 5% horse serum for 2 hours and incubated overnight with the primary antibody at 4 °C. The following day, sections were incubated with DAPI (1:10,000) for 5 min followed three time of washing with PBS. Whole-mount tissues and cryo-sections were recorded with a Zeiss LSM 880 confocal laser scanning microscope with ZEN microscopy software, and image analysis was processed by Imaris software (Oxford instruments).

For spatial analysis, the position of each cell was determined based on the thresholds for the fluorescence intensity of its soma using the spot detection function in IMARIS. Cell identification was then performed using a Boolean gating strategy based on the different fluorescence signals, as shown graphically in the figure. The average distances to the nearest TdTom^+^ Iba1^+^ cells were used to color-code clustering behavior. Border-corrected kernel density estimations of TdTom^+^Iba1^+^ cells were calculated in R studio as previously described ^60^ and displayed using the *ggplot2* package. The following primary antibodies were used: rabbit polyclonal anti-Iba1 (1:250, Wako, cat. no. 019-19741, RRID: AB_839504), rat monoclonal anti-CD31 (1:250, MEC13.3, Biolegend, cat. no. 102502, RRID: AB_312909), rabbit monoclonal anti-Tmem119 (1:200, 28-3, Abcam, cat. no. ab209064, RRID: AB_2800343), rat monoclonal anti-Lyve1 conjugated with eFluor660 (1:200, ALY7, Invitrogen, cat. no. 50-0443-82, RRID: AB_10597449), goat polyclonal anti-MMR/CD206 (1:250, R&D systems, cat. no. AF2535, RRID: AB_2063012), rabbit polyclonal anti-Tyrosine hydroxylase (1:400, Merk Millipore, cat. no. AB152, RRID: AB_390204), and rabbit monoclonal anti-MHC Class II (I-A/I-E) (1:250, M5/114.15.2, eBioscience, cat. no. 14-5321-82, RRID: AB_467561). Donkey anti-rat IgG (H+L) Cy2 (cat. no. 712-225-153), donkey anti-rat IgG (H+L) cy3 (cat. no. 712-165-154), donkey anti-rabbit IgG (H+L) Cy2 (cat no. 711-225-152), and donkey anti-goat IgG (H+L) Alexa Fluor 647 (cat. no. 705-605-147) were used for secondary antibodies from Jackson Immuno Research Laboratory (JIR).

### RNAseq libraries preparation

For the bulk-RNA sequencing (RNAseq) library preparation, 10^5^ or less (when isolated cell number was insufficient) microglia and leptomeningeal MΦ were sorted into 40 µL of lysis/binding buffer (Life Technologies) and stored at −80 °C until mRNA isolation. mRNA was captured with Dynabeads oligo(dT) (Life Technologies) according to the manufacturer’s guidelines. A derivation of MARSseq protocol ^78^ was used to prepare libraries for RNAseq. For the single-cell RNA sequencing (scRNAseq) library preparation, cells are sorted in a 384-well plate with lysis buffer and poly-T barcoded primers and placed in −80C for future processing. When thawed, they are heated to bind the primer to the mRNA and first strand mix is added for cDNA-mRNA hybrid. ExoI is added to remove excess primers and all samples are then pooled followed by second-strand synthesis. Pooled samples are amplified overnight using IVT, proceeded by RNA fragmentation and ligation of adapter for pooled sample identification. Finally, first strand synthesis and PCR for final sample amplification. Library concentration was measured with a Qubit fluorometer (Life Technologies) and mean molecule size was determined with a 2200 TapeStation instrument. RNaseq libraries were sequenced using the Illumina NextSeq 500 with 90 reads, 75 for read 1, which contained the ligation adapter sequence and mRNA sequence, and 15 for read 2 containing the single sample or single cell barcode and UMI sequences.

### RNAseq analysis

Bulk-RNAseq data analysis was performed by using the UTAP transcriptome analysis pipeline ^79^. Gene expression levels were calculated and normalized using DESeq2 with the parameters: betaPrior = True, cooksCutoff = FALSE, independent Filtering = FALSE. Batch correction was done using the sva (3.26.0) R when batch adjustments were required. Raw p-values were adjusted for multiple testing, using the procedure of Benjamini-Hochberg method. Differentially expressed genes were selected with absolute fold change >= 2, and adjusted p-value =<0.05. The gene expression matrix was clustered using the *k*-means algorithm. Heatmaps were generated using Morpheus (Broad Institute). For the scRNAseq data analysis, sequence mapping and read count performed using the MARS-seq pipeline ^78^.

### MERFISH spatial transcriptomics

In order to map major cell types and statement of the cells, a MERFISH gene panel for 140 genes was designed. The target gene probe library was constructed by Vizgen. Mice were anesthetized with Pental (1:1 with PBS) and perfused with PBS. Brains were isolated and fixed with 4% paraformaldehyde in room temperature for 2 hours, and transferred to 30% sucrose solution which is 4% paraformaldehyde based. 30% sucrose incubation has done for 48 hours at 4 °C, and tissues were imbedded in optimal-cutting-temperature compound (TissueTek) for frozen blocks. Tissues were cryosectioned at 10 µm for sagittal section to expose both striatum and substantia nigra. Two sagittal sections were mount onto 40 mm cover slips (Vizgen supplied) coated with poly-ethyleneimine, and dried for 15 min at 4 °C. Sections were washed 3 times in PBS for 5 min and dehydrated in 70% ethanol. Tissue-mounted coverslips were processed according to Vizgen’s protocol and imaged on a MERSCOPE alpha scope (Vizgen), imaging seven z-planes with DAPI, poly T, and GFP sequential panel detection. Each MERFISH round consisted of readout probe hybridization (10 min), washing (5 min), and imaging of ∼1,000 tiled fields of view (FOV) (220 μm x 220 μm per FOV), readout fluorophore cleavage by TCEP (15 min), and rinsing with 2 x SSC (5 min). For each round, images were acquired with 750-nm, 650-nm, and 560-nm lasers at six focal planes separated by 1 μm in z to image the readout probes. Images of DAPI and poly T in sequential panels were used to cell segmentation as demonstrated previously ^80^. Using a deep learning-based cell segmentation algorithm (Cellpose) ^81^, central position of each segmented nucleus in each z-plane was identified, and connected across different z-plane. Cell soma were segmented by poly T RNA image also with Cellpose algorithm. Together with segmented nuclei and segmented cell bodies each cell segmentation was determined. Encoded MERFISH images of RNA spots were de-multiplexed using MERlin (https://github.com/emanuega/MERlin), using the codebook provided by Vizgen. The MERFISH count matrices were processed replicating the procedure described in ^80^. Gene counts for each cell were normalized by the total count per cell. Data dimensionality was reduced with PCA, followed by identification of cell clusters, and visualized by UMAP similar to the approach described for scRNAseq data analysis with the Jupiter code that provided by Vizgen, and visualization of transcriptomes on single cells was performed by MERSCOPE Vizualizer (Vizgen).

### Behavior tests

Before tests all animals was placed in a reverse light cycle room for 2 weeks. The assessment of gait was performed with the CatWalk XT system. Each rodent could explore and walk freely through the alley without any rewards. No dedicated session for habituation to the CatWalk apparatus was performed. The CatWalk software automatically recorded the video of each run, where the rodent covered the whole distance of the walkway, and total number of records in each time point was 3 with cut off of maximum variation <= 50%. An experimental session lasts for 5–15 minutes. The positions of the footprints in the recorded videos were identified automatically by the software and then corrected manually by an experienced observer. For the pole test, each mouse was placed head-up on top of a vertical wooden pole with a rough surface, 50 cm in height and 1 cm in diameter. The animals were allowed to orient themselves downward and to descend along the pole back into their home cage. Each mouse was exposed to three trials, and the time spent to orient downward (t-turn) and the time to descend (t-descend) were recorded.

### Quantification and statistical analysis

In all experiments, individual values are presented with mean and standard deviation (SD). Statistical tests were selected based on appropriate assumptions with respect to data distribution and variance characteristics. Student’s t test (two-tailed) was applied to demonstrate statistical differences between two groups, and multiple t test (two-tailed) was performed for comparisons between two groups with different time points. Statistical significance was defined as *p<0.05*, and symbols mean *p < 0.0001* (****), *p < 0.001* (***), *p < 0.01* (**), and *p < 0.05* (*). Otherwise, specific *p values* are written. Sample sizes were chosen according to standard guidelines. Number of animals is indicated as “n” and presented with the number of dots on the graphs. Animals of the same age, sex, and genetic background were randomly assigned to treatment groups.

## Online Supplement

**Supplementary Movie 1.**

Cryo-sagittal section of 6-month-old *Ms4a3^cre/+^: R26^TdTomato^* mouse brain stained with CD31 antibodies. Enlarged images come from the green rectangles in the sagittal section image. Scale is indicated in the sagittal section image, and scale bar of enlarged images is 15 µm. (see also **Suppl. Fig 1d**).

**Supplementary Movie 2.**

Merscope analysis of sagittal brain sections using Merfish gene list in Suppl. Data 1.

## Supplementary Figure legends

**Supplementary Figure 1.**
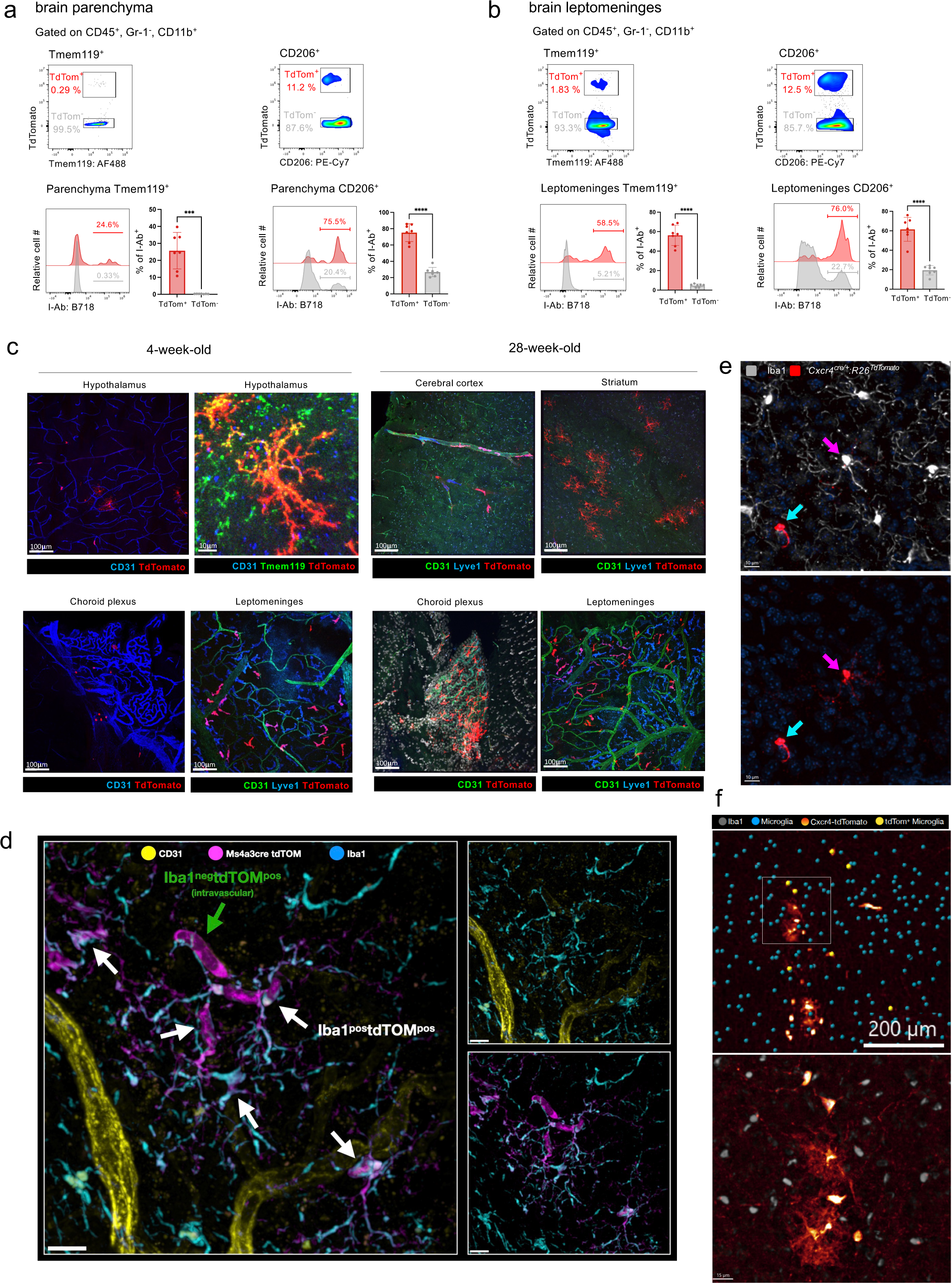
Flow cytometry and microscope images of brains of *Ms4a3^cre/+^: R26^TdTomato^* mice. (a) Representative flow cytometry results of parenchymal brain cells of *Ms4a3^cre/+^: R26^TdTomato^* mice comparing MHCII expression levels between MoMg and microglia. Pre-gated on CD45^+^ and Ly6C/G^−^. (n = 7). (b) Representative flow cytometry results of leptomeningeal brain cells of *Ms4a3^cre/+^: R26^TdTomato^* mice comparing MHCII expression levels between MoMΦ and endogenous MΦ. Pre-gated on CD45^+^ and Ly6C/G^−^. (n = 7). (c) Representative images of 4- and 28-week-old *Ms4a3^cre/+^: R26^TdTomato^* mouse brains. Scale is indicated in each image. (d) Representative image of cryo-sagittal section of 6-month-old *Ms4a3^cre/+^: R26^TdTomato^*mouse brain stained with CD31 antibodies. Scale is indicated in the sagittal section image, and scale bar of enlarged images is 15 µm. (see also Supplementary movie 1). (e) Representative image of cryo-sagittal brain section 6-month-old *Cxcr4^creer/+^: R26^TdTomato/TdTomato^* mouse that was TAM pulsed at 6 weeks of age and sacrificed 4 months later for analysis, stained with Iba1 antibodies. Note presence of ramified MoMg. Scale bar is 10 µm. (f) Representative image of MoMΦ with microglia morphology residing in diencephalon/midbrain region of 6-month-old *Cxcr4^creer/+^: R26^TdTomato/TdTomato^* mouse that was TAM pulsed at 6 weeks of age and sacrificed 4 months later for analysis, stained with Iba1 antibodies. Scale bars indicated.

**Supplementary Figure 2.**
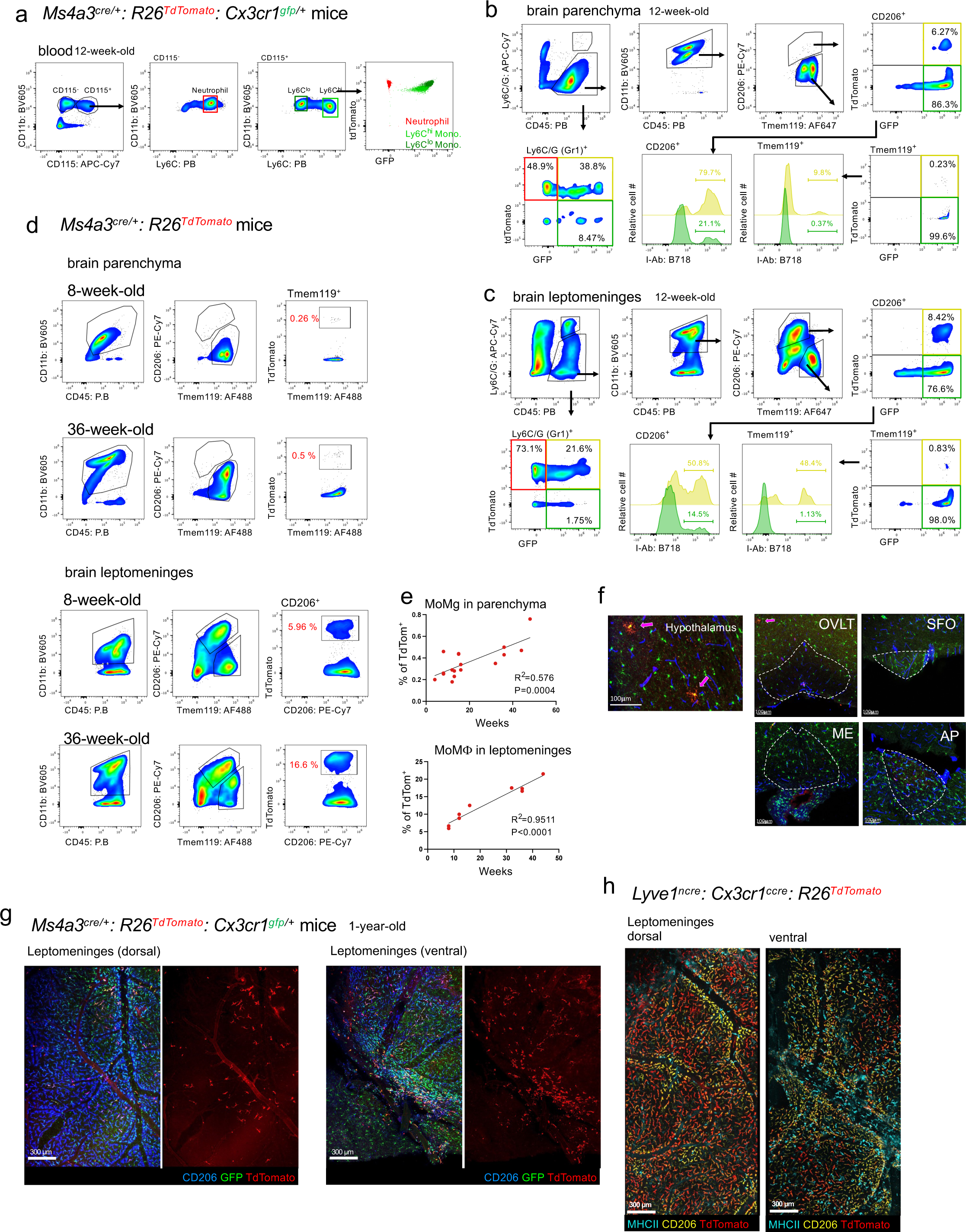
Flow cytometry and microscope images of brains from *Ms4a3^cre/+^: R26^TdTomato^:Cx3cr1^gfp/+^*, *Ms4a3^cre/+^: R26^TdTomato^* and *Lyve1^ncre^: Cx3cr1^ccre^: R26^TdTomato^* mice. (a) Flow cytometer plots of *Ms4a3^cre/+^: R26^TdTomato^: Cx3cr1^gfp/+^* mouse blood. Neutrophils are defined as CD115^−^, Ly6C^int^. (b) Representative flow cytometry results of brain parenchymal cells of *Ms4a3^cre/+^: R26^TdTomato^: Cx3cr1^gfp/+^* mouse brain and histogram to show MHCII positive cell percentages between endogenous and monocyte-derived cells. (c) Representative flow cytometry results of brain leptomeningeal cells of *Ms4a3^cre/+^: R26^TdTomato^: Cx3cr1^gfp/+^* mouse brain and histogram to show MHCII positive cell percentages between endogenous and monocyte-derived cells. (d) Representative flow cytometry results of brain parenchymal and leptomeningeal cells of 8- and 36-week-old *Ms4a3^cre/+^: R26^TdTomato^* mice brains to show the difference of monocyte-derived cell fractions. (e) Correlation analysis of monocyte-derived cell and mouse ages in both parenchymal Tmem119^+^ and leptomeningeal CD206^+^ populations in *Ms4a3^cre/+^: R26^TdTomato^* mouse brains. (f) Representative images of circumventricular organs (CVO) of 20-week-old *Ms4a3^cre/+^: R26^TdTomato^: Cx3cr1^gfp/+^* mouse. GFP^+^ cells indicate YS-derived microglia, and GFP^+^, TdTomato^+^ cells indicate MoMg. Magenta arrows indicates GFP^+^, TdTomato^+^ cells. *organum vasculosum of the lamina terminalis* (OVLT), *subfornical organ* (SFO), *median eminence* (ME), and *area postrema* (AP). (g) Tile scan images of ventral and dorsal leptomeninges from 1-year-old *Ms4a3^cre/+^: R26^TdTomato^: Cx3cr1^gfp/+^* mouse brain. Whole-mount samples were stained with anti CD206 antibodies. Scale is indicated in each image. (h) Tile scan images of ventral and dorsal leptomeninges from 5-month-old *Lyve1^ncre^: Cx3cr1^ccre^: R26^TdTomato^* mouse brain. Whole-mount samples were stained with anti MHCII and anti CD206 antibodies. Scale is indicated in each image.

**Supplementary Figure 3.**
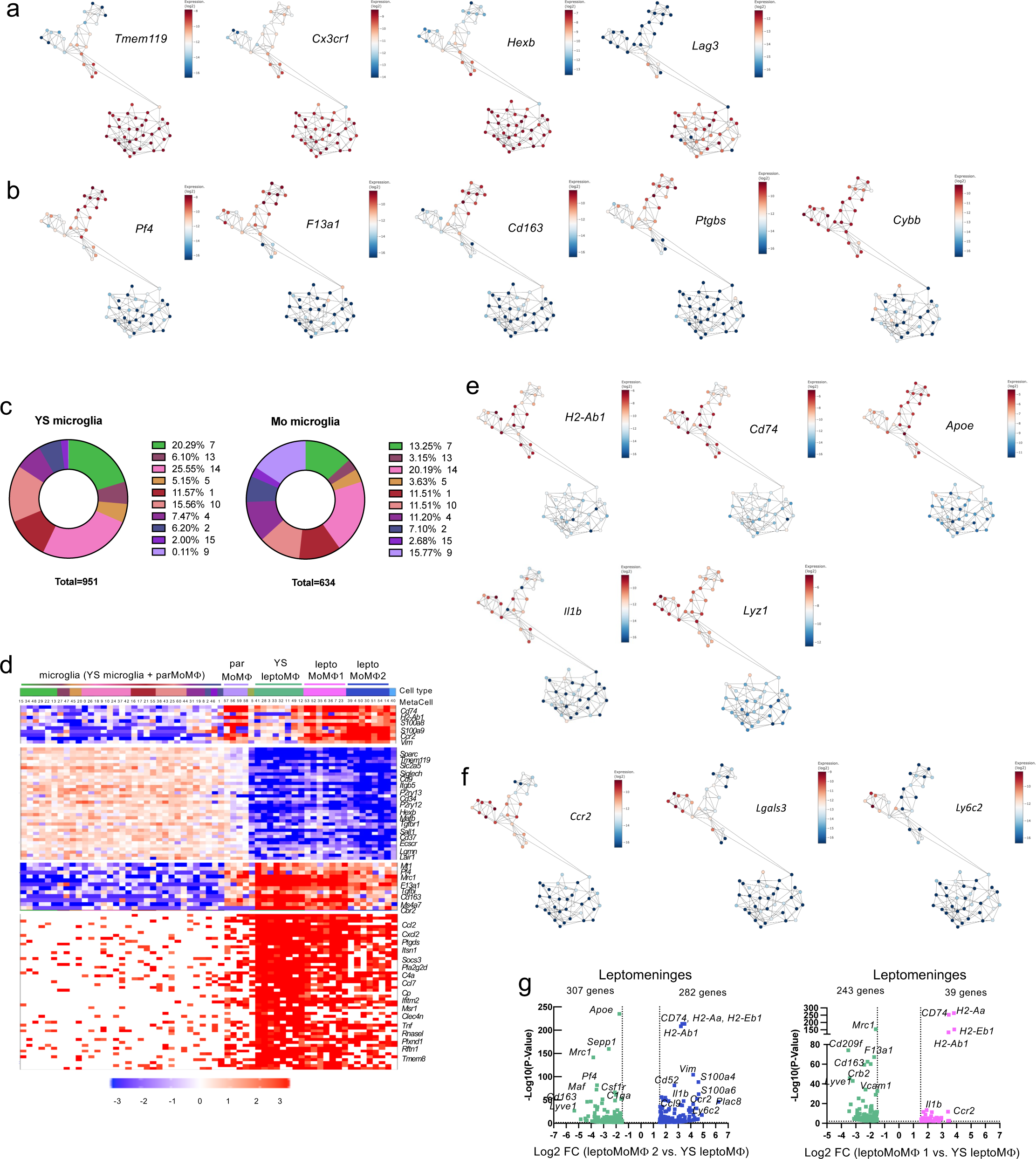
Single cell RNA sequencing of parenchymal and leptomeningeal MΦ from *Ms4a3^cre/+^: R26^TdTomato^: Cx3cr1^gfp/+^* mice. (a) MetaCell plots showing gene expression levels of microglia markers between the MetaCells. (b) MetaCell plots showing gene expression levels of leptomeningeal MΦ markers between the MetaCells. (c) Pie graphs showing number of sorted cells yielding MetaCells. (d) Heatmap showing distinctive gene module expression between different Meta cell types. Gene modules are separated with space. (e) MetaCell plots showing representative gene expression levels for MHCII and activation. (f) MetaCell plots showing gene expression level related with monocyte-derived cells. (g) Volcano plots showing the difference between the different leptomeningeal MΦ types. Different colors are used for each cell clusters, and representative genes are indicated. Only significant DEG are plotted. Significance means -Log10 (p value) > 1.5, Log 2 (fold change) > 1.

**Supplementary Figure 4.**
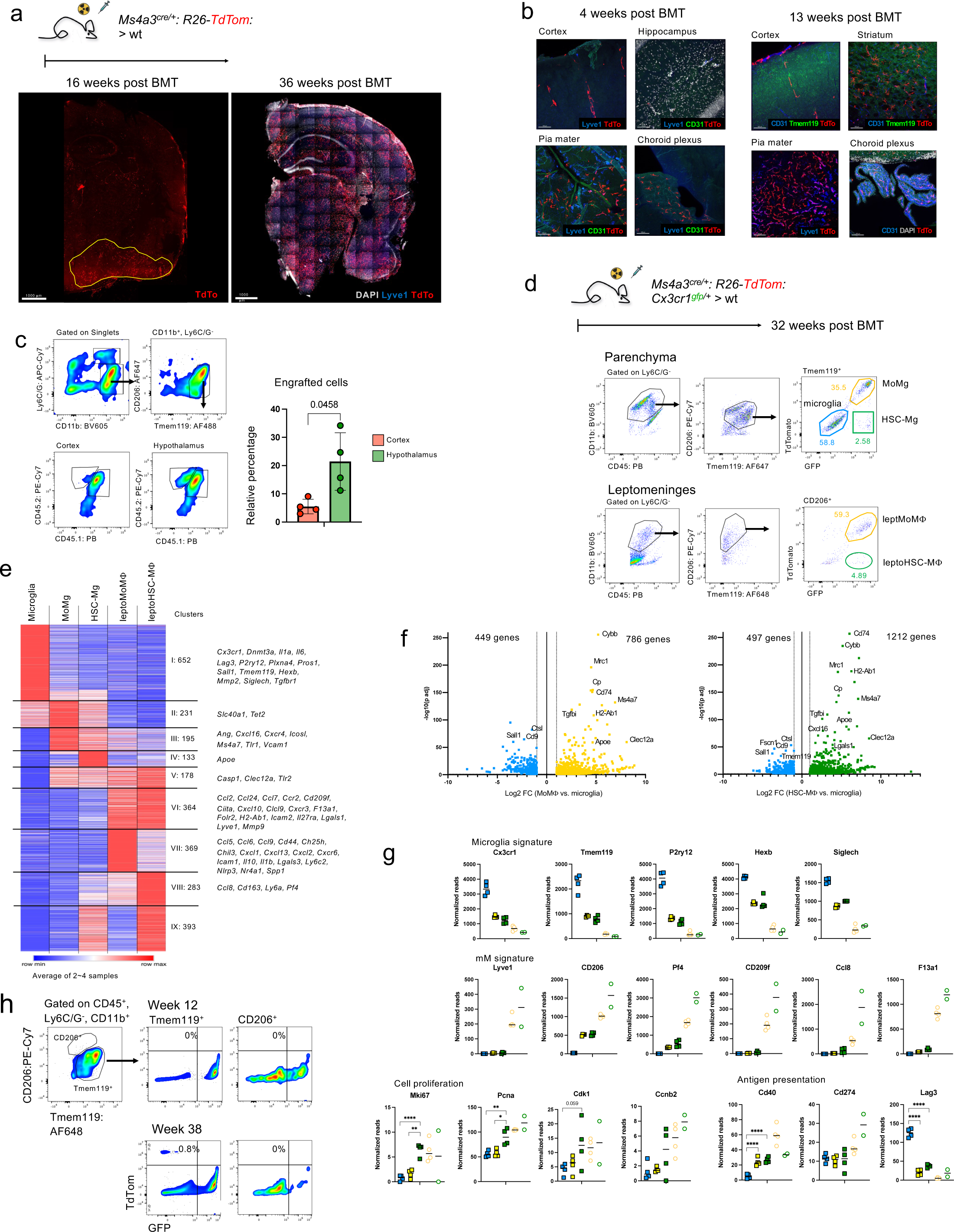
Flow cytometry, histology and bulk RNA sequencing results of brains of BM chimeras. (a) BMT scheme with *Ms4a3^cre/+^: R26^TdTomato^* BM transferred to lethally irradiated WT mice. Coronal section images of brains, 16 and 36 weeks after BMT showing the cell infiltration pattern. Regions where ramified TdTomato^+^ cells accumulated 16 weeks post BMT are indicated with a yellow line. Scales are indicated in each image. (b) Representative images of brain regions, 4 and 13 weeks after BMT. Bar scale represents 100 µm. (c) Flow cytometric quantification of MoMg in cortex and hypothalamus of old CD45.2 > CD45.1 BM chimeras. Representative flow cytometry blot, left; bar graph summary, right, (n = 4). (d) BMT scheme with *Ms4a3^cre/+^: R26^TdTomato^: Cx3cr1^gfp/+^* BM transferred into lethally irradiated WT mice and flow cytometry plots for cell sorting of each population. (e) Heatmap of transcriptomes of indicated sorted cell populations. (f) Volcano plots of transcriptomes comparing microglia and MoMg, and microglia and HSC-MΦ. Only significant DEG are plotted, and significance means -Log10 (adj. p value) > 1.5, Log 2 (fold change) > 1 (g) Normalized gene expression of selected genes. (n = 4 or 2). (h) Representative flow cytometry plots of parenchymal brain cells of chimeras generated with *Sall1^ncre^: Cx3cr1^ccre^: R26^TdTomato^* BM transferred into irradiated *Cx3cr1^gfp/+^* mice, 12 and 38 weeks after BMT. Plots are pre-gated on CD45^+^, Ly6C/G^−^, and CD11b^+^ cells.

**Supplementary Figure 5.**
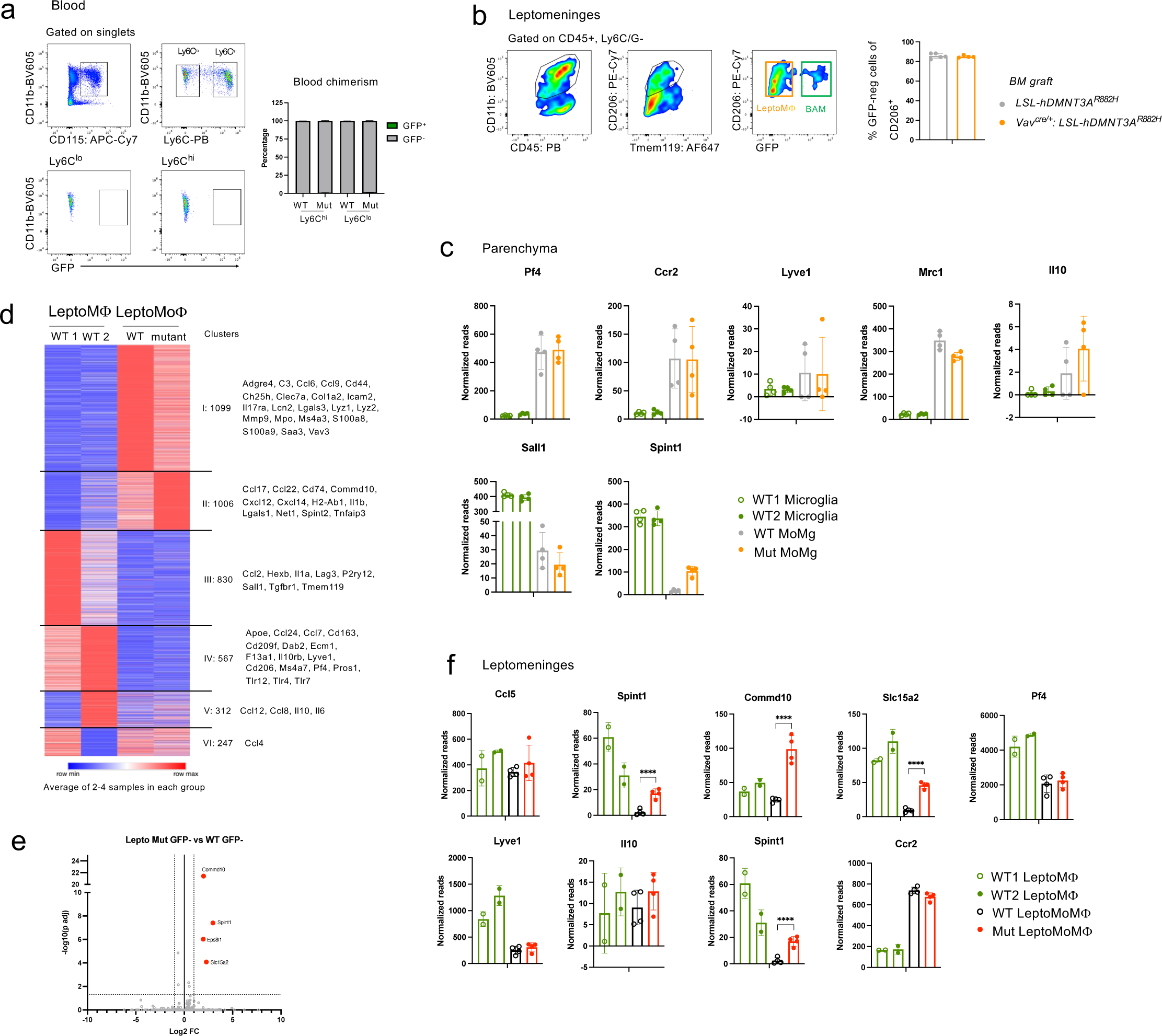
Flow cytometry and bulk RNA sequencing results of *Vav^cre/+^: hDNMT3A^R822H/+^* BM chimeras and controls. (a) Flow cytometry result of blood from Cx3cr1^gfp/+^ mice that *Vav^cre/+^: hDNMT3A^R822H/+^* BM transferred after lethal dose irradiation. n = 5. (b) Representative flow cytometry plots of brain leptomeningeal cells, pre-gated for CD45 and Ly6C/G, 18 weeks after BMT. Bar graph shows leptomeningeal cell chimerism of mutated and control groups. (n = 5). (c) Normalized reads of selected genes differentially expressed between control and mutated engrafted cells from brain parenchyma. (n = 4). (d) Heatmap of bulk RNA sequencing result from leptomeningeal cell compartment. Average values of 2-4 samples. (e) Volcano plot comparing control, MoMΦ and mutated MoMΦ in the leptomeninges of brains of the chimeras. (f) Normalized reads of selected genes differentially expressed between control and mutated engrafted cells isolated brain leptomeninges. (n = 4 or 2).

**Supplementary Figure 6.**
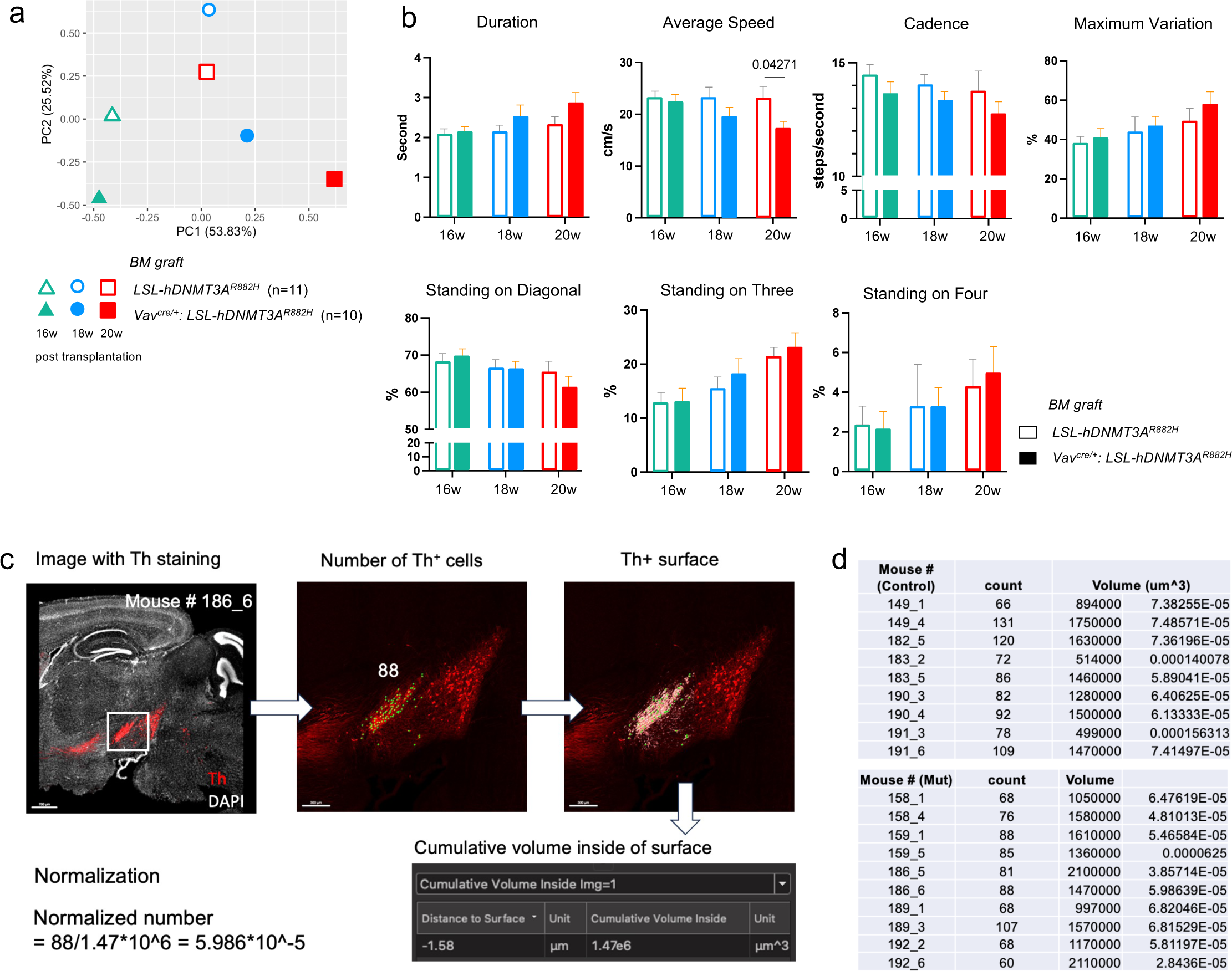
Analysis of Gait and quantification of dopaminergic neurons in substantia nigra of *Vav^cre/+^: hDNMT3A^R822H/+^* BM chimeras and controls. (a) PCA plots of 120 parameters acquired with CatWalk gait analysis system. Time points of CatWalk analysis are 16, 18 and 20 weeks after BMT; experimental repeat with independently generated groups of BM chimeras. (n=10, per group). (b) CatWalk gait results of parameters that indicated motor impairment of mutated BM recipients compared with the control BM recipients. Multiple T-test was performed for statistical comparison, q values are written in the graphs. (n = 10, per group). (c) Representative image of TH staining of sagittal section of brain of recipient of mutated BM and illustration of approach for quantification according to Th^+^ areas, as determined automatically by Imaris. (d) Summary of counts of TH^+^ neurons in *substantia nigra pars compacta* of recipients of control and mutant BM.

**Supplementary Table 1.** Quantification of microglia and MoMg in selected brain regions on sagittal brain sections of young and old *Ms4a3^cre/+^: R26^TdTomato^: Cx3cr1^gfp/+^*mouse (related to Fig. 2E). Mg are defined as being Cx3cr1-GFP positive, MoMg are Cx3cr1-GFP positive and TdTom-positive.

|  |  | young |  |  | mean | old |  |  | mean |
| --- | --- | --- | --- | --- | --- | --- | --- | --- | --- |
| Frontal cortex | Mg | 3797 | 3952 | 4107 |  | 4572 | 4185 | 4727 |  |
|  | MoMg | 0 | 0 | 0 |  | 0 | 1 | 1 |  |
|  | Ratio (%) | 0.00 | 0.00 | 0.00 | **0** | 0.00 | 0.02 | 0.02 | **0.015** |
| Midbrain | Mg | 1700 | 1232 | 1232 |  | 1742 | 1360 | 1530 |  |
|  | MoMg | 9 | 3 | 8 |  | 9 | 7 | 16 |  |
|  | Ratio (%) | 0.53 | 0.24 | 0.65 | **0.47** | 0.52 | 0.51 | 1.05 | **0.695** |
| Diencephalon | Mg | 4070 | 4347 | 3145 |  | 4440 | 3607 | 4717 |  |
|  | MoMg | 11 | 16 | 19 |  | 47 | 39 | 41 |  |
|  | Ratio (%) | 0.27 | 0.37 | 0.60 | **0.41** | 1.06 | 1.08 | 0.87 | **1.003** |
| Subst.nigra | Mg | 209 | 198 | 143 |  | 231 | 209 | 220 |  |
|  | MoMg | 4 | 2 | 2 |  | 2 | 9 | 6 |  |
|  | Ratio (%) | 1.91 | 1.01 | 1.39 | **1.44** | 0.86 | 4.29 | 2.72 | **2.625** |

## References

1. Prinz, M., Jung, S. & Priller, J. Microglia Biology: One Century of Evolving Concepts. Cell 179, 292–311 (2019).

2. Ratz, M. et al. Clonal relations in the mouse brain revealed by single-cell and spatial transcriptomics. Nat Neurosci 25, 285–294 (2022).

3. Ginhoux, F. et al. Fate Mapping Analysis Reveals That Adult Microglia Derive from Primitive Macrophages. Science 330, 841–845 (2010).

4. Askew, K. et al. Coupled Proliferation and Apoptosis Maintain the Rapid Turnover of Microglia in the Adult Brain. CellReports 18, 391–405 (2017).

5. Tay, T. L. et al. A new fate mapping system reveals context-dependent random or clonal expansion of microglia. Nature Neuroscience 20, 793–803 (2017).

6. Varol, C., Mildner, A. & Jung, S. Macrophages: development and tissue specialization. Annual Review of Immunology 33, 643–675 (2015).

7. Goldmann, T. et al. Origin, fate and dynamics of macrophages at central nervous system interfaces. Nat Immunol 17, 797–805 (2016).

8. Perdiguero, E. G. et al. Tissue-resident macrophages originate from yolk-sac-derived erythro-myeloid progenitors. Nature 1–17 (2014) doi:10.1038/nature13989.

9. Goldmann, T. et al. A new type of microglia gene targeting shows TAK1 to be pivotal in CNS autoimmune inflammation. Nat Neurosci 16, 1618–1626 (2013).

10. Brioschi, S. et al. A Cre-deleter specific for embryo-derived brain macrophages reveals distinct features of microglia and border macrophages. Immunity 56, 1027–1045.e8 (2023).

11. Shechter, R. et al. Infiltrating blood-derived macrophages are vital cells playing an anti-inflammatory role in recovery from spinal cord injury in mice. PLoS medicine 6, e1000113 (2009).

12. Ajami, B., Bennett, J. L., Krieger, C., Mcnagny, K. M. & Rossi, F. M. V. Infiltrating monocytes trigger EAE progression, but do not contribute to the resident microglia pool. Nature Neuroscience 14, 1142–1149 (2011).

13. Ajami, B., Bennett, J. L., Krieger, C., Tetzlaff, W. & Rossi, F. M. V. Local self-renewal can sustain CNS microglia maintenance and function throughout adult life. Nature Neuroscience 10, 1538–1543 (2007).

14. Shemer, A. et al. Engrafted parenchymal brain macrophages differ from microglia in transcriptome, chromatin landscape and response to challenge. Nat Commun 9, 5206 (2018).

15. Bennett, F. C. et al. A Combination of Ontogeny and CNS Environment Establishes Microglial Identity. Neuron 98, 1170–1183.e8 (2018).

16. Cronk, J. C. et al. Peripherally derived macrophages can engraft the brain independent of irradiation and maintain an identity distinct from microglia. The Journal of experimental medicine 47, jem.20180247–21 (2018).

17. Lund, H. et al. Competitive repopulation of an empty microglial niche yields functionally distinct subsets of microglia-like cells. Nat Commun 9, 1–13 (2018).

18. Fixsen, B. R. et al. SALL1 enforces microglia-specific DNA binding and function of SMADs to establish microglia identity. Nat. Immunol. 1–12 (2023) doi:10.1038/s41590-023-01528-8.

19. Buttgereit, A. et al. Sall1 is a transcriptional regulator defining microglia identity and function. Nat Immunol 17, 1397–1406 (2016).

20. Wolf, Y. et al. Microglial MHC class II is dispensable for experimental autoimmune encephalomyelitis and cuprizone-induced demyelination. Eur. J. Immunol. 48, 1308–1318 (2018).

21. Giladi, A. et al. Cxcl10+ monocytes define a pathogenic subset in the central nervous system during autoimmune neuroinflammation. Nat Immunol 21, 525–534 (2020).

22. Amorim, A. et al. IFNγ and GM-CSF control complementary differentiation programs in the monocyte-to-phagocyte transition during neuroinflammation. Nat Immunol 23, 217–228 (2022).

23. Wicks, E. E. et al. The Translational Potential of Microglia and Monocyte-Derived Macrophages in Ischemic Stroke. Front Immunol 13, 897022 (2022).

24. Werner, Y. et al. Cxcr4 distinguishes HSC-derived monocytes from microglia and reveals monocyte immune responses to experimental stroke. Nat Neurosci 23, 351–362 (2020).

25. ElAli, A. & Rivest, S. Microglia in Alzheimer’s disease: A multifaceted relationship. Brain, Behavior, and Immunity 55, 138–150 (2016).

26. Yan, P. et al. Peripheral monocyte-derived cells counter amyloid plaque pathogenesis in a mouse model of Alzheimer’s disease. J. Clin. Investig. 132, e152565 (2022).

27. Spiteri, A. G., Wishart, C. L., Pamphlett, R., Locatelli, G. & King, N. J. C. Microglia and monocytes in inflammatory CNS disease: integrating phenotype and function. Acta Neuropathol. 143, 179–224 (2022).

28. Bouzid, H. et al. Clonal hematopoiesis is associated with protection from Alzheimer’s disease. Nat. Med. 1–9 (2023) doi:10.1038/s41591-023-02397-2.

29. Shlush, L. I. Age-related clonal hematopoiesis. Blood 131, 496–504 (2018).

30. Jaiswal, S. & Ebert, B. L. Clonal hematopoiesis in human aging and disease. Science 366, (2019).

31. Fuster, J. J. et al. Clonal hematopoiesis associated with Tet2 deficiency accelerates atherosclerosis development in mice. Science eaag1381–11 (2017) doi:10.1126/science.aag1381.

32. Wang, W. et al. Macrophage Inflammation, Erythrophagocytosis, and Accelerated Atherosclerosis in Jak2V617F Mice. Circ Res 123, e35–e47 (2018).

33. Sano, S. et al. CRISPR-Mediated Gene Editing to Assess the Roles of Tet2 and Dnmt3a in Clonal Hematopoiesis and Cardiovascular Disease. Circ Res 123, 335–341 (2018).

34. Evans, M. A., Sano, S. & Walsh, K. Cardiovascular Disease, Aging, and Clonal Hematopoiesis. Annu. Rev. Pathol.: Mech. Dis. 15, 1–20 (2019).

35. Alliot, F., Godin, I. & Pessac, B. Microglia derive from progenitors, originating from the yolk sac, and which proliferate in the brain. Brain research. Developmental brain research 117, 145–152 (1999).

36. Schulz, C. et al. A lineage of myeloid cells independent of Myb and hematopoietic stem cells. Science 336, 86–90 (2012).

37. Kierdorf, K. et al. Microglia emerge from erythromyeloid precursors via Pu.1- and Irf8-dependent pathways. Nature Neuroscience (2013) doi:10.1038/nn.3318.

38. Liu, Z. et al. Fate Mapping via Ms4a3-Expression History Traces Monocyte-Derived Cells. Cell 178, 1509–1525.e19 (2019).

39. Hove, H. V. et al. A single-cell atlas of mouse brain macrophages reveals unique transcriptional identities shaped by ontogeny and tissue environment. Nature Neuroscience 22, 1021–1035 (2019).

40. Jung, S. et al. Analysis of Fractalkine Receptor CX3CR1 Function by Targeted Deletion and Green Fluorescent Protein Reporter Gene Insertion. Mol Cell Biol 20, 4106–4114 (2000).

41. Kim, J.-S. et al. A Binary Cre Transgenic Approach Dissects Microglia and CNS Border-Associated Macrophages. Immunity 54, 176–190.e7 (2021).

42. Amit, I., Winter, D. R. & Jung, S. The role of the local environment and epigenetics in shaping macrophage identity and their effect on tissue homeostasis. Nature Immunology 17, 18–25 (2016).

43. Baran, Y. et al. MetaCell: analysis of single-cell RNA-seq data using K-nn graph partitions. 1–19 (2019) doi:10.1186/s13059-019-1812-2.

44. Zhang, X. et al. DNMT3A and TET2 compete and cooperate to repress lineage-specific transcription factors in hematopoietic stem cells. Nat Genet 48, 1014–1023 (2016).

45. Xie, M. et al. Age-related mutations associated with clonal hematopoietic expansion and malignancies. Nat Med 20, 1472–1478 (2014).

46. Fabre, M. A. et al. The longitudinal dynamics and natural history of clonal haematopoiesis. Nature 606, 335–342 (2022).

47. Abelson, S. et al. Prediction of acute myeloid leukaemia risk in healthy individuals. Nature 559, 400–404 (2018).

48. Cobo, I. et al. DNA methyltransferase 3 alpha and TET methylcytosine dioxygenase 2 restrain mitochondrial DNA-mediated interferon signaling in macrophages. Immunity (2022) doi:10.1016/j.immuni.2022.06.022.

49. Scheller, M. et al. Hotspot DNMT3A mutations in clonal hematopoiesis and acute myeloid leukemia sensitize cells to azacytidine via viral mimicry response. Nat Cancer 2, 527–544 (2021).

50. Zioni, N. et al. Inflammatory signals from fatty bone marrow support DNMT3A driven clonal hematopoiesis. Nat. Commun. 14, 2070 (2023).

51. Russler-Germain, D. A. et al. The R882H DNMT3A Mutation Associated with AML Dominantly Inhibits Wild-Type DNMT3A by Blocking Its Ability to Form Active Tetramers. Cancer Cell 25, 442–454 (2014).

52. Kim, S. J. et al. A DNMT3A mutation common in AML exhibits dominant-negative effects in murine ES cells. Blood 122, 4086–4089 (2013).

53. Festa, B. P. et al. Microglial-to-neuronal CCR5 signaling regulates autophagy in neurodegeneration. Neuron (2023) doi:10.1016/j.neuron.2023.04.006.

54. Ajoy, R. et al. CCL5 promotion of bioenergy metabolism is crucial for hippocampal synapse complex and memory formation. Mol Psychiatr 26, 6451–6468 (2021).

55. Chou, S.-Y. et al. CCL5/RANTES contributes to hypothalamic insulin signaling for systemic insulin responsiveness through CCR5. Sci Rep-uk 6, 37659 (2016).

56. Timotius, I. K. et al. Systematic data analysis and data mining in CatWalk gait analysis by heat mapping exemplified in rodent models for neurodegenerative diseases. J Neurosci Meth 326, 108367 (2019).

57. Matsuura, K., Kabuto, H., Makino, H. & Ogawa, N. Pole test is a useful method for evaluating the mouse movement disorder caused by striatal dopamine depletion. J Neurosci Meth 73, 45–48 (1997).

58. Tandon, R., Yadav, G., Singh, B. P. & Gupta, A. K. Gait analysis of patients with Parkinson-plus syndromes: a research article. Bull. Natl. Res. Cent. 47, 76 (2023).

59. Yamasaki, R. et al. Differential roles of microglia and monocytes in the inflamed central nervous system. Journal of Experimental Medicine 211, 1533–1549 (2014).

60. Silvin, A. et al. Dual ontogeny of disease-associated microglia and disease inflammatory macrophages in aging and neurodegeneration. Immunity 55, 1448–1465.e6 (2022).

61. Xiong, B. et al. Precise Cerebral Vascular Atlas in Stereotaxic Coordinates of Whole Mouse Brain. Front Neuroanat 11, 128 (2017).

62. Blinder, P. et al. The cortical angiome: an interconnected vascular network with noncolumnar patterns of blood flow. Nature Neuroscience 16, 889–897 (2013).

63. Davalos, D. et al. Fibrinogen-induced perivascular microglial clustering is required for the development of axonal damage in neuroinflammation. Nature Communications 3, 1227 (2012).

64. Shemer, A. et al. Interleukin-10 Prevents Pathological Microglia Hyperactivation following Peripheral Endotoxin Challenge. Immunity 53, 1033–1049.e7 (2020).

65. Skelly, D. T., Hennessy, E., Dansereau, M.-A. & Cunningham, C. A Systematic Analysis of the Peripheral and CNS Effects of Systemic LPS, IL-1Β, TNF-α and IL-6 Challenges in C57BL/6 Mice. Plos One 8, e69123–20 (2013).

66. Keogh, M. J. et al. High prevalence of focal and multi-focal somatic genetic variants in the human brain. Nat Commun 9, 4257 (2018).

67. Huarte, O. U. et al. Single-Cell Transcriptomics and In Situ Morphological Analyses Reveal Microglia Heterogeneity Across the Nigrostriatal Pathway. Front. Immunol. 12, 639613 (2021).

68. Moca, E. N. et al. Microglia Drive Pockets of Neuroinflammation in Middle Age. J. Neurosci. 42, 3896–3918 (2022).

69. Masuda, T. et al. Specification of CNS macrophage subsets occurs postnatally in defined niches. Nature 604, 740–748 (2022).

70. Siddhartha, J. et al. Age-Related Clonal Hematopoiesis Associated with Adverse Outcomes. New Engl J Med 371, 2488–2498 (2014).

71. Arends, C. M. et al. Associations of clonal hematopoiesis with recurrent vascular events and death in patients with incident ischemic stroke. Blood 141, 787–799 (2022).

72. Poewe, W. et al. Parkinson disease. Nat Rev Dis Primers 3, 17013 (2017).

73. Zimprich, A. et al. Mutations in LRRK2 Cause Autosomal-Dominant Parkinsonism with Pleomorphic Pathology. Neuron 44, 601–607 (2004).

74. Bick, A. G. et al. Genetic IL-6 Signaling Deficiency Attenuates Cardiovascular Risk in Clonal Hematopoiesis. Circulation 141, 124–131 (2019).

75. Madisen, L. et al. A robust and high-throughput Cre reporting and characterization system for the whole mouse brain. Nat Neurosci 13, 133–140 (2009).

76. Stadtfeld, M. & Graf, T. Assessing the role of hematopoietic plasticity for endothelial and hepatocyte development by non-invasive lineage tracing. Development 132, 203–213 (2004).

77. Cossarizza, A. et al. Guidelines for the use of flow cytometry and cell sorting in immunological studies (second edition). Eur J Immunol 49, 1457–1973 (2019).

78. Keren-Shaul, H. et al. MARS-seq2.0: an experimental and analytical pipeline for indexed sorting combined with single-cell RNA sequencing. Nature Protocols 1–25 (2019) doi:10.1038/s41596-019-0164-4.

79. Kohen, R. et al. UTAP: User-friendly Transcriptome Analysis Pipeline. BMC bioinformatics 20, 154–7 (2019).

80. Moffitt, J. R. et al. Molecular, spatial, and functional single-cell profiling of the hypothalamic preoptic region. Science 362, (2018).

81. Stringer, C., Wang, T., Michaelos, M. & Pachitariu, M. Cellpose: a generalist algorithm for cellular segmentation. Nat. Methods 18, 100–106 (2021).

